# Pathogenic mitochondrial DNA mutations inhibit melanoma metastasis

**DOI:** 10.1101/2023.09.01.555986

**Authors:** Spencer D. Shelton, Sara House, Vijayashree Ramesh, Zhenkang Chen, Tao Wei, Xun Wang, Claire B. Llamas, Siva Sai Krishna Venigalla, Cameron J. Menezes, Zhiyu Zhao, Jennifer G. Gill, Ralph J. DeBerardinis, Sean J. Morrison, Alpaslan Tasdogan, Prashant Mishra

## Abstract

Mitochondrial DNA (mtDNA) mutations are frequently observed in cancer, but their contribution to tumor progression is controversial. To evaluate the impact of mtDNA variants on tumor growth and metastasis, we created human melanoma cytoplasmic hybrid (cybrid) cell lines transplanted with wildtype mtDNA or pathogenic mtDNA encoding variants that partially or completely inhibit oxidative phosphorylation. Homoplasmic pathogenic mtDNA cybrids reliably established tumors despite dysfunctional oxidative phosphorylation. However, pathogenic mtDNA variants disrupted spontaneous metastasis of subcutaneous tumors and decreased the abundance of circulating melanoma cells in the blood. Pathogenic mtDNA did not induce anoikis or inhibit organ colonization of melanoma cells following intravenous injections. Instead, migration and invasion were reduced, indicating that limited circulation entry functions as a metastatic bottleneck amidst mtDNA dysfunction. Furthermore, analysis of selective pressure exerted on the mitochondrial genomes of heteroplasmic cybrid lines revealed a suppression of pathogenic mtDNA allelic frequency during melanoma growth. Collectively, these findings demonstrate that functional mtDNA is favored during melanoma growth and enables metastatic entry into the blood.

## Introduction

Pathogenic mutations within the mitochondrial genome (mtDNA) are widely recognized as causative for inherited diseases, yet their role in the pathology of acquired diseases is largely unknown^1-3^. Somatic mtDNA mutations commonly occur in human tumors, with an incidence rate greater than 50%^4-6^. Certain tumor types such as colorectal, thyroid, and renal cancers exhibit a disproportionately high incidence and allelic burden of deleterious mtDNA mutations^5-9^, and are typically associated with “oncocytic” changes secondary to excessive mitochondrial accumulation^10^. However, high allelic deleterious mtDNA mutations are atypical in most tumors, and the majority of cancers maintain somatic mtDNA mutations at a low allelic frequency^4-6^. Additionally, most cancer types appear to favor functional mtDNA, as indicated by a lower allelic frequency of deleterious mutations relative to nonpathogenic variants^4-6^. These correlation studies have yet to be functionally investigated in an experimental model that rigorously evaluates the impact of mtDNA variants on tumor progression in otherwise isogenic backgrounds. In particular, *in vivo* studies with nuclear isogenic tumors are needed to test whether selective pressures maintain functioning mtDNA in most tumor types.

A number of studies have suggested that healthy mitochondria promote cancer progression. For example, disseminated cancer cells in melanoma, breast, and renal cancers display increased expression of the mitochondrial biogenesis transcription factor PGC-1α, which promotes mitochondrial mass and oxygen consumption^11-13^. In oral squamous cell carcinoma, metastatic cells are observed to enhance mtDNA translation through mt-tRNA modifications^14^. From a metabolic standpoint, activity in the tricarboxylic acid (TCA) cycle is elevated in metastatic tumors of triple-negative breast cancer and clear cell renal cell carcinoma when compared to primary tumors^15,16^. Further, pharmacologic inhibition of mitochondrial complex I with IACS-010759 in pre-clinical models curbed melanoma brain metastases without significantly impeding primary tumor growth^17^. While these findings indicate that metastatic dissemination stimulates mitochondrial activities, it has been reported that variants in the mitochondrial genome can convey metastatic potential^18-20^. Clarifying how the mitochondrial genome influences the metastatic cascade could reconcile the role for mtDNA in metastasis and highlight the potential for mtDNA targeted anti-cancer treatments.

Engineering genetically encoded mitochondrial dysfunction within a respiring and innately metastatic tumor model would enable direct elucidation of the specific metastatic processes predominantly dependent upon mitochondrial activities. However, it has been reported that ablation of subunits of complex I, II, and III obstructs tumor growth through mechanisms not associated with oxidative phosphorylation, but reliant on ubiquinol oxidation^21^. Consequently, creation of models that only partially compromise mitochondrial function may bypass these tumor growth inhibitory effects. Moreover, the consequences of mutations in complex IV and V remain uncharacterized in the context of tumor progression.

In this study, we addressed the consequences of loss-of-function mtDNA mutations in tumor metastasis utilizing the endogenously metastatic human melanoma cell line A375 as a model system. We developed a flow cytometry-based protocol to transplant wildtype or patient-derived dysfunctional mitochondrial genomes into the A375 cell line, thereby creating nuclear-isogenic cell lines with distinct mtDNA variants. Investigated pathogenic mtDNA variants resulted in either partial or complete inhibition of mitochondrial electron transport chain (ETC) function. Despite mtDNA-encoded functional deficits, all homoplasmic cybrid models invariably established tumors. These mtDNA mutant tumor models demonstrated a pronounced reduction in spontaneous metastasis, primarily attributed to a diminished potential for tumor cells to infiltrate the bloodstream. Furthermore, we assessed selective pressures exerted on the mitochondrial genome of heteroplasmic cybrid lines, revealing that melanoma growth selects against pathogenic mtDNA variants. Cumulatively, these findings provide the first direct *in vivo* experimental verification of mtDNA selection during melanoma growth and demonstrate that functional mtDNA promotes metastatic entry into the blood.

## Results

### Generation of melanoma cybrid models with loss-of-function mtDNA variants

In the immortalized human melanoma cell line A375, we established isogenic cytoplasmic hybrid (cybrid) models each harboring distinct mitochondrial genomes. For cybrid model generation, the endogenous mtDNA needed to be depleted so that exogenous mtDNA sources could repopulate the mtDNA pool. Following a two weeks treatment with 5 μM or 10 μM dideoxycytidine (ddC), an irreversible inhibitor of mtDNA replication^22^, we established multiple A375 clones in which mtDNA was reduced to undetectable levels (Extended Data Fig. 1a). While the parental line demonstrated functional mitochondrial oxygen consumption, clones from both ddC treatment concentrations exhibited no mitochondrial oxygen consumption (Extended Data Fig. 1b,c). For cybrid generation, clones treated with 5 μM ddC were selected as the mtDNA-depleted (ρ0) recipient line.

Existing cybrid fusion protocols rely on antibiotic and metabolic selection to eliminate unfused contaminating donor cells and ρ0 acceptor cells^23,24^. To enhance the efficiency of cybrid generation and render the protocol more versatile across cell lines, we integrated cellular compartment staining followed by flow cytometry-based enrichment, effectively removing the necessity for antibiotic and metabolic selection (Extended Data Fig. 2a). Initially, the mitochondria and nuclei of mtDNA donor cells were stained with MitoTracker Green (MTGreen) and Hoechst 33342. Stained donor cells were then incubated with cytochalasin B, an actin polymerization inhibitor, and subjected to a high speed centrifugation atop a percoll cushion, thereby generating a mixture of nuclei negative cytoplasts and whole cells^25^ (Extended Data Fig. 2b). Enucleated cytoplasts, identified as the Hoechst^−^,MTGreen^+^ population, were enriched through flow cytometry (Extended Data Fig. 2c). These enriched cytoplasts were subsequently fused with A375 ρ0 cells that had been pre-stained with the nuclear dye SYTO59 (Extended Data Fig. 2a). Finally, fused cybrid cells, identified as the SYTO59^+^,MTGreen^+^,Hoechst^−^ population, were enriched by flow cytometry (Extended Data Fig. 2d).

To evaluate the impact of mtDNA variants on tumor growth and progression, we generated a panel of A375 homoplasmic cybrids (variant allele frequency (VAF) = 1) (Fig. 1a). These cybrids carried either wildtype (WT) mtDNA (with no pathogenic variants) or pathogenic variants associated with human disease (Table 1). We established and validated multiple independent clonal lines for each cybrid model, ensuring the retention of the A375 nuclear genome (through STR analysis) and successful transplantation of the mtDNA genome by Sanger sequencing (Extended Data Fig. 3a-c). To precisely verify homoplasmic allelic frequency for the partial loss of function models, ATP6 and ND1, we utilized digital droplet PCR (ddPCR) (Fig. 1b,c, Extended Data Fig. 3d-i). The pathologic deletion in the complete loss of function model, CO1, involves a frameshift within a homopolymeric region, preventing quantitative ddPCR analysis. However, western blot analysis revealed a loss of mt.CO1 protein expression, while expression of another mtDNA encoded protein, mt.ATP8, remained intact (Fig. 1d).

**Figure 1.**
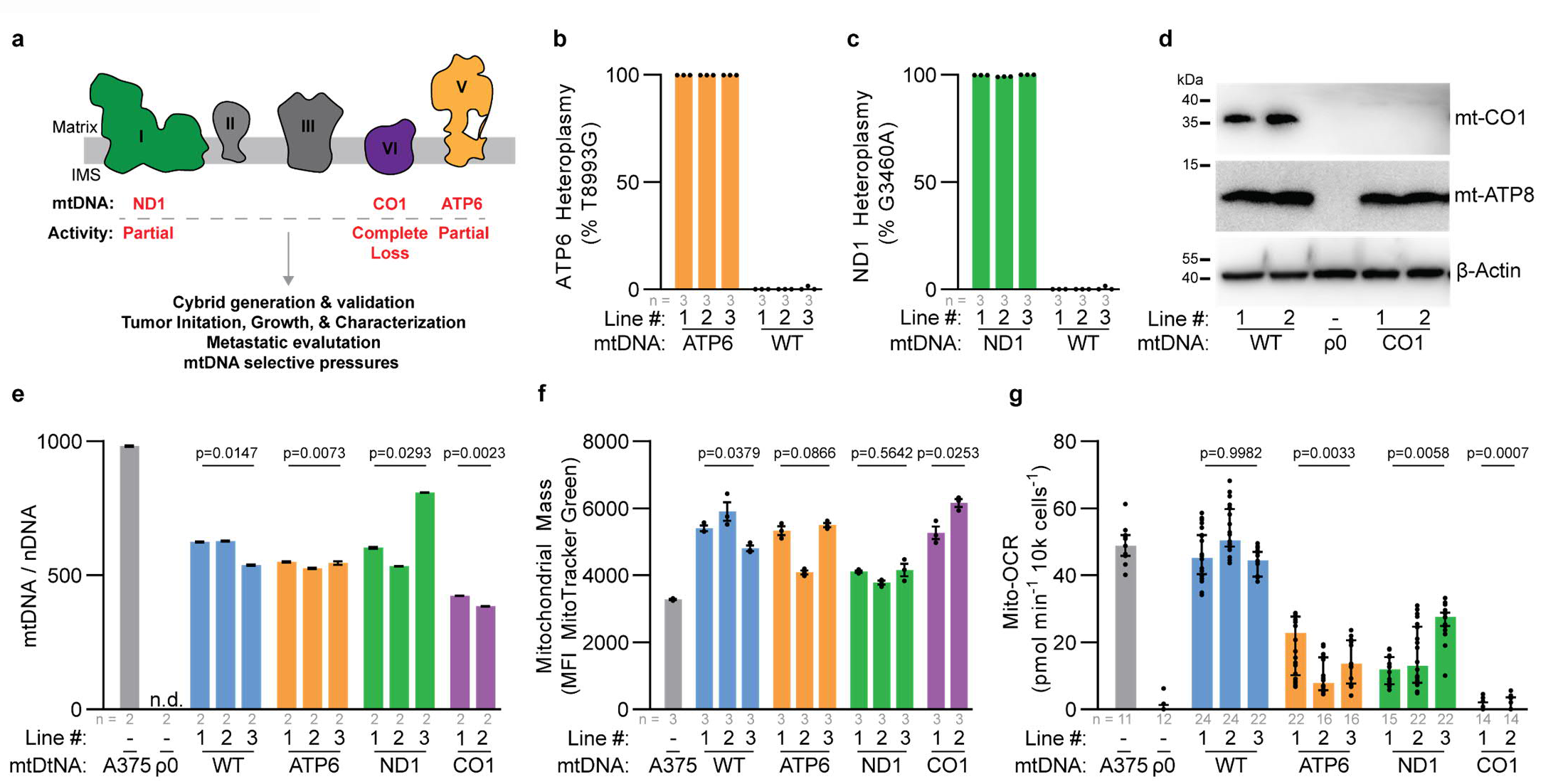
mtDNA pathogenic variants in A375 homoplasmic cybrid lines impact mitochondria ETC activity. **a**, Schematic of electron transport chain with complexes labeled to display subunits with mtDNA variants investigated in this study. **b**,**c**, ddPCR quantitation of allelic fraction for (**b**) ATP6 allele mt.T8993G and (**c**) ND1 allele mt.G3460G in homoplasmic cybrid lines. **d**, Western blot analysis for the indicated mtDNA encoded proteins in homoplasmic cybrid cell lines. β-Actin levels are shown as loading control. **e**, Mitochondrial genome (mtDNA) to nuclear genome (nDNA) ratios in the indicated homoplasmic cybrid lines. n.d., not detected. P values reflect comparisons with the A375 parental line. **f**, Mitochondrial mass as assessed by flow cytometric analysis of MitoTracker Green staining in the indicated homoplasmic cybrid lines. P values reflect comparisons with the A375 parental line. MFI, mean fluorescence intensity. **g**, Mitochondrial oxygen consumption rates (Mito-OCR) in indicated homoplasmic cybrid lines. P values reflect comparisons with the A375 parental line. The number of samples analyzed per treatment is indicated. Data are mean (**b**,**c**), mean ± s.e.m. (**e**,**f**), and median ± interquartile range (**g**). Statistical significance was assessed using one-way ANOVA with Dunn’s multiple comparison adjustment (**d**) and nested one-way ANOVA with Dunn’s multiple comparison adjustment (**f**,**g**).

**Table 1.** Origin of cybrid donor mtDNA.

| **Cybrid** | **ETC Complex** | **NT Change (AA Change)** | **Oxidative Activity** | **Origin of mtDNA** |
| --- | --- | --- | --- | --- |
| **WT** | - | - | Full Activity | Verified healthy donor |
| **ATP6** | V | T8993G (L156R) | Partial Loss | mtDNA disease Maternally Inherited Leigh Syndrome (MILS) & Neuropathy Ataxia and Retinitis Pigmentosa (NARP) |
| **ND1** | I | G3460A (A52T) | Partial Loss | mtDNA disease Leber Hereditary Optic Neuropathy (LHON) |
| **CO1** | IV | 6692.Del (Truncation) | Complete Loss | Melanoma Patient Derived Xenograft |

All cybrid lines displayed a restoration of mitochondrial genome content relative to the A375 ρ0 clone, albeit at levels lower than the parental line (Fig. 1e). The mitochondrial mass (based on MTGreen staining) of the cybrid lines was generally elevated relative to the parental line (Fig. 1f). The influence of the mitochondrial genome on oxygen consumption reflected the anticipated functional consequences of the respective pathogenic mtDNA alleles^26,27^. Specifically, the mitochondrial oxygen consumption rate was unchanged between the parental line and the WT cybrids, partially reduced in the ATP6 and ND1 cybrids, and completely ablated in the CO1 cybrids (Fig. 1g).

### Tumor growth is sustained in mtDNA dysfunctional cybrids

We subcutaneously xenografted the A375 cybrids into the hind flank of immunocompromised NOD.CB17-Prkdc^scid^ Il2rg^tm1Wjl^/SzJ (NSG) mice and monitored tumor growth over time (Fig. 2a). Despite variations in ETC capacity, all cybrid models reliably established tumors at either 100 or 10,000 cell injections (Fig. 2b-c). Upon tumor harvesting, we confirmed the *in vivo* stability of the homoplasmic pathogenic mtDNA variants (Extended Data Fig. 4a-c). Observed growth rates among the subcutaneous tumors were heterogenous, with a general reduction in comparison to the parental line – an effect further accentuated in the models with mtDNA dysfunction (Fig. 2c). Assessment of Ki67 staining across the tumors revealed comparable levels of proliferation amongst the cybrid models (Fig. 2d, Extended Data Fig. 5a-d). Histological analysis indicated substantial areas of tumor necrosis within the WT, ND1, and ATP6 cybrid tumors (Fig. 2e,f, Extended Data Fig. 6a). Conversely, the CO1 tumors exhibited negligible tumor necrosis and displayed an increased prevalence of disorganized, or discohesive, regions (Fig. 2e,f, Extended Data Fig. 6a-c).

**Figure 2.**
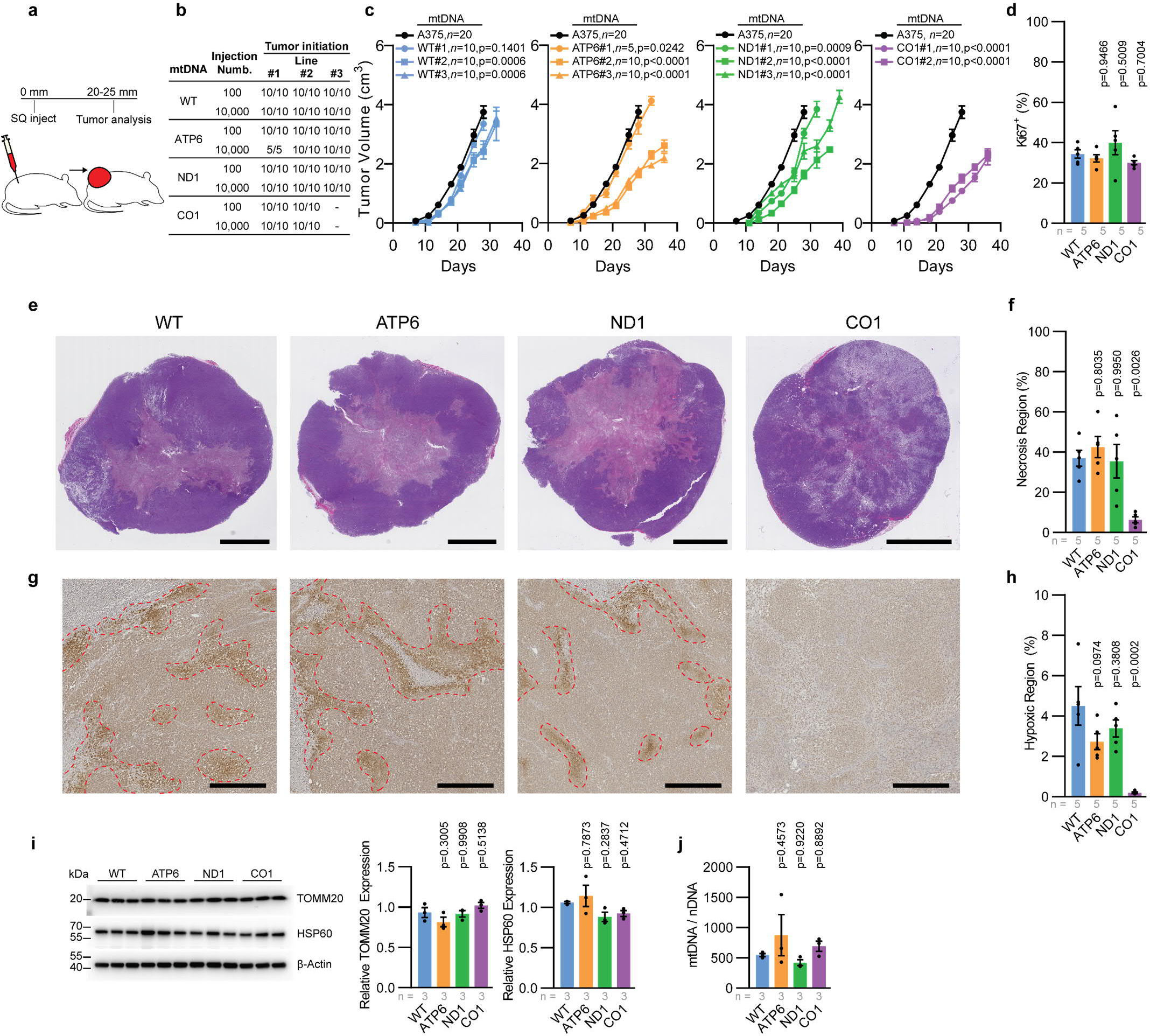
Functional mtDNA is dispensable for primary melanoma growth. **a**, Illustration of subcutaneous injection to assess primary tumors of homoplasmic cybrid lines. **b**, Frequency of tumor formation for indicated injection counts across homoplasmic cybrid lines. **c**, Tumor growth rate of indicated cybrid lines following 10,000 cell subcutaneous injection. Homoplasmic growth data (black circle) is repeated as a reference in all panels. P values reflect comparison with wildtype (WT) group. **d**, Quantitation of the percentage of Ki67^+^ nuclei across subcutaneous tumors. P values reflect comparison with wildtype (WT) group. **e**, Representative H&E images of subcutaneous tumors. Scale bar, 5000 μm. **f**, Quantitation of necrotic region as a percentage of tumor cross-sectional area. P values reflect comparison with wildtype (WT) group. **g**, Representative immunohistochemistry images for pimonidazole from subcutaneous tumors of the indicated mtDNA haplotype. Scale bar, 500 μm. **h,** Quantitation of pimonidazole positive (hypoxic) regions as a percentage of tumor cross-sectional area. P values reflect comparison with wildtype (WT) group. **i**, Western blot analysis of mitochondrial outer membrane protein TOMM20 and matrix protein HSP60. β-Actin expression is shown as loading control. P values reflect comparison with wildtype (WT) group. **j**, Mitochondrial genome content (mtDNA/nDNA) for subcutaneous tumors of the indicated mtDNA haplotype. The number of samples analyzed per group is indicated. Data are mean ± s.e.m. (**d**,**f**,**h**,**i**,**j**). Statistical significance was assessed using exponential growth least squares fitting on the mean values of replicates followed by extra sum-of-squares F test with Holm-Sidak’s adjustment (**b**) and one-way ANOVA with Dunn’s multiple comparison adjustment (**d**,**f**,**h**,**i**,**j**).

Pimonidazole staining revealed that tumors with WT, ATP6, and ND1 mtDNA contained comparable levels of hypoxia (Fig. 2g,h). In contrast, no detectable hypoxic regions were observed in tumors with the CO1 variant (Fig. 2g,h). Interestingly, we failed to find significant differences in mitochondrial biomass, assessed by measuring mitochondrial protein expression (TOMM20 on the outer membrane, and HSP60 in the matrix), and mitochondrial genome content among the cybrid tumors (Fig. 2i,j), indicating a lack of oncocytic transformation. Collectively, the pathogenic mtDNA variants did not preclude the growth of subcutaneous melanoma xenografts, as evidenced by 100% of implants forming tumors, but generally reduced tumor growth rates. While the WT, ATP6, and ND1 presented with comparable tumor morphology, the most severe loss of function model, CO1, presented with histological, hypoxic, and necrotic variations.

### Pathogenic mtDNA variants suppress spontaneous metastasis

The xenografted lines were engineered to express luciferase, enabling quantitation of spontaneous metastatic disease burden via bioluminescence imaging of organs. Specifically, the total spontaneous metastatic burden, as measured by bioluminescence of dissected organs, was analyzed when primary subcutaneous tumors attained a size of 20-25 mm in diameter (Fig. 3a). Both the WT and partial loss of function ATP6 cybrid tumors exhibited spontaneous metastasis at levels comparable to the parental line (Fig. 3b,c). In contrast, the ND1 cybrid tumors exhibited a substantial decrease in spontaneous metastatic burden, and no metastasis was detected from the CO1 cybrid tumors (Fig. 3b,c). Correspondingly, there was a substantial reduction in the frequency of circulating melanoma cells in the blood of all mtDNA mutant cybrid lines (Fig. 3d, Extended Data Fig. 7a,b).

**Figure 3.**
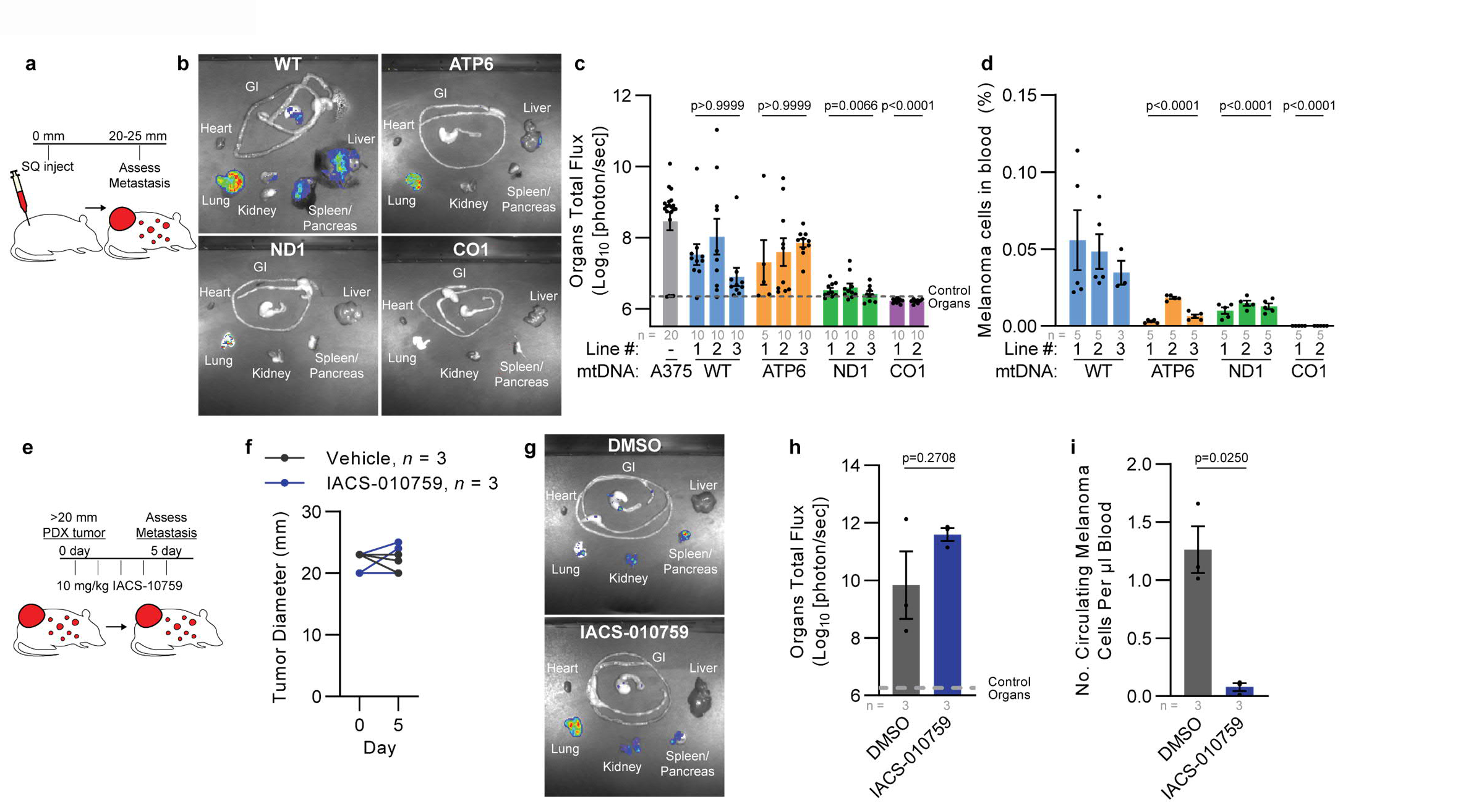
Spontaneous metastasis is reduced by dysfunctional mtDNA. **a**, Illustration of subcutaneous injection to assess spontaneous metastasis. **b**,**c**, Representative images (**b**) and quantitation (**c**) of metastatic bioluminescence signal of organs from mice with late-stage subcutaneous xenografts of indicated cybrid lines. P values indicate comparisons with the A375 (parental) group. **d**, Frequency of circulating melanoma cells from mice (as a percentage of DAPI negative events) with late-stage subcutaneous xenografts of indicated cybrid lines. P values indicate comparisons with WT group. **e**, Illustration of acute pharmacological evaluation of IACS-010759 to assess metastasis in late-stage patient derived xenograft model UT10. **f**, Tumor diameter of xenografted UT10 mice treated with IACS-010759 or vehicle. **g**,**h**, Representative images (**g**) and quantitation (**h**) of organ bioluminescence signal of UT10 xenograft following treatment with IACS-010759 or vehicle. **i**, Circulating melanoma cells normalized to total blood volume of xenografted UT10 mice following treatment with IACS-010759 or vehicle. The number of tumors or mice analyzed per treatment is indicated. Data are mean ± s.e.m. (**c**,**d**,**h**,**i**). Statistical significance was assessed using non-parametric Kruskal-Wallis test with Dunn’s multiple comparison adjustment (**c**), nested one-way ANOVA with Dunn’s multiple comparison adjustment (**d**), two-tailed unpaired t-test with Welch’s correction (**h,i**).

To extend these findings, we investigated whether pharmacologic inhibition of mitochondrial ETC function in tumors with functional mtDNA suppresses the emergence of circulating melanoma cells in the blood. Mice bearing advanced-stage melanoma patient-derived xenograft (PDX) UT10^28,29^ tumors were subjected to an acute 5-day oral gavage of either 10 mg/kg IACS-010759 (an established bioavailable complex I inhibitor^30^) or 0.5% methylcellulose vehicle control (Fig. 3e). The short-term IACS-010759 treatment did not induce changes in primary tumor size or organ metastatic burden (Fig. 3f-h). However, IACS-010759 treatment led to a significant decrease in the number of circulating melanoma cells in the blood (Fig. 3i). These findings indicate that either genetic or pharmacologic impairment of mitochondrial ETC activity can inhibit the appearance of melanoma cells in the blood.

### Pathogenic mtDNA variants inhibit tumor cell motility and invasion

Considering that the onset of anoikis may limit metastasis^31^, we investigated whether the cybrid models exhibited differential detachment survival potentials. In line with observations of metabolic perturbation induced by detachment^32-35^, detached culture of the A375 cybrid models increased reactive oxygen species (ROS), reduced glucose consumption, and reduced lactate excretion (Fig. 4a-c). Relative to the WT line, these effects were exacerbated in the complete loss of function CO1 model. Despite these detachment induced stresses, the mtDNA mutant cybrid lines exhibited significantly elevated cell counts following 24 hours of detached culture relative to the WT line (Fig. 4d). Detachment resulted in only marginal reductions in viability and minimal increases in apoptosis for all cybrid lines (Fig. 4e,f). To directly investigate metastatic seeding from the blood stream, we injected 1,000 cells of each cybrid line into the tail vein of NSG mice (Fig. 4g). Live BLI imaging showed significant bioluminescence signal in all cybrid lines, irrespective of ETC capacity (Fig. 4h). Further analysis of dissected organs indicated no significant difference in the total metastatic disease burden (Fig. 4i). These results demonstrate that mtDNA mutations do not significantly inhibit the ability of melanoma cells to survive detachment or seed metastatic sites following direct bloodstream injections.

**Figure 4.**
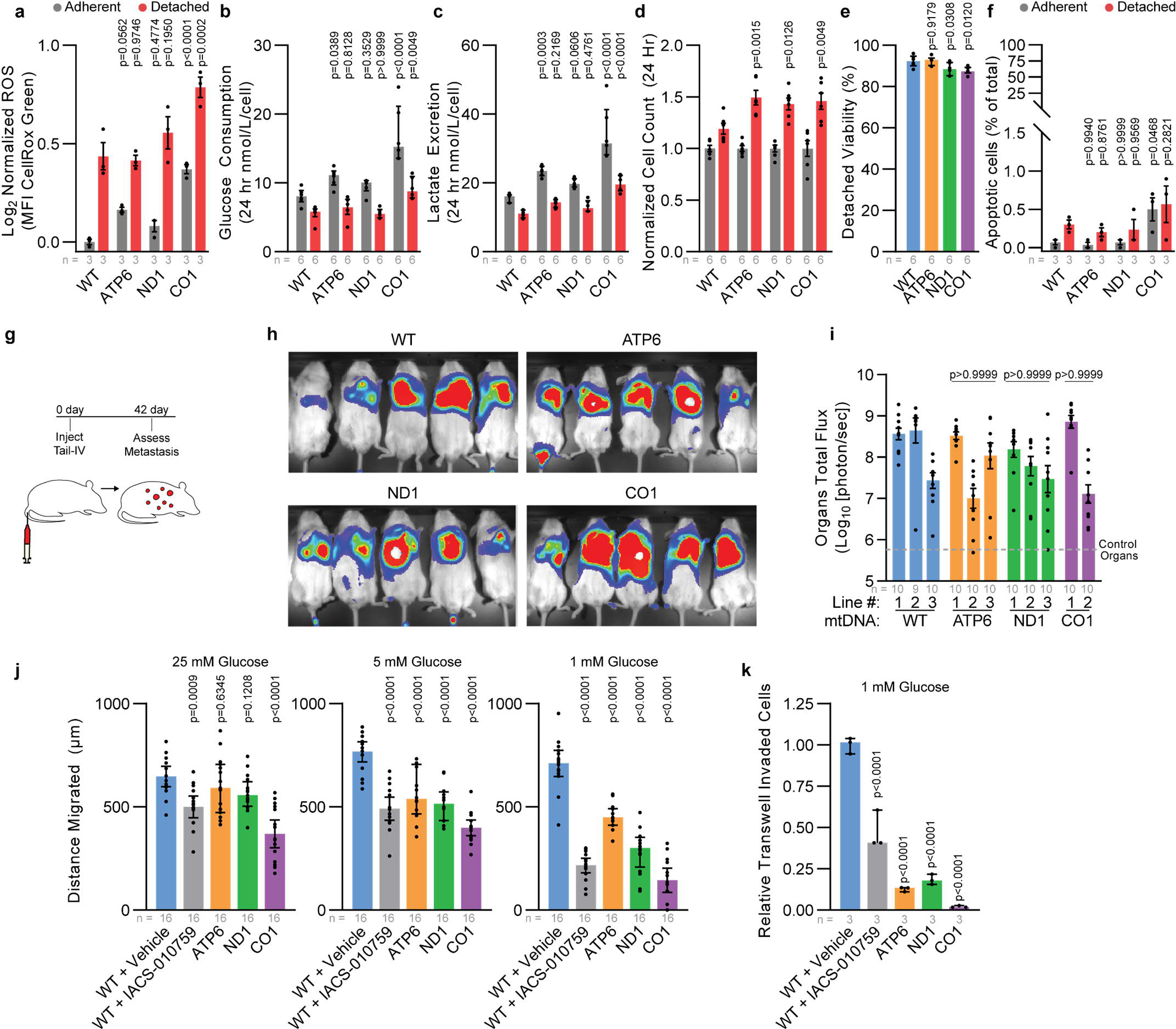
Dysfunctional mtDNA disrupts migratory potential in human melanoma. **a-d**, Reactive oxygen species (**a**), glucose consumption (**b**), lactate excretion (**c**), and cell count (**d**) for indicated cybrid cells after 24 hour adherent and detached culture. P values indicate comparison with WT group for each condition. MFI, mean fluorescence intensity **e**, Detached viability of indicated cybrid lines following 24 hour of culture. P values indicate comparison with WT group. **f**, Percentage of apoptotic cells following 24 hour culture in adherent or detached conditions. P values indicate comparison with WT group. **g**, Illustration of intravenous injection of cybrid lines to assess organ colonization. **h**,**i**, Representative bioluminescence imaging of live mice (**h**) and quantitation of metastatic bioluminescence signal of organs (**i**) following intravenous injection of indicated cybrid lines. P values indicate comparisons with WT group. **j**, 24 hour wound healing assay quantification for indicated cybrid lines at 25 mM, 5 mM, and 1 mM glucose concentrations. Gap distance was quantified from the difference of the 0 hour and 24 hour wound gap. P values indicate comparisons with WT + vehicle group for each glucose concentration. **k,** Relative transwell invaded cells at 1 mM glucose concentration. P values indicate comparisons with WT + vehicle group for each glucose concentration. The number of samples, tumors, or mice analyzed per treatment is indicated. Data are mean ± s.e.m. (**a**), median ± interquartile range (**i**,**j**,**k**). Statistical significance was assessed using one-way ANOVA with Dunn’s multiple comparison adjustment (**e**), non-parametric Kruskal-Wallis test with Dunn’s multiple comparison adjustment (**i**), two-way ANOVA with Dunn’s multiple comparison adjustment (**a**,**b**,**c**,**d**,**f**), and one-way ANOVA with Dunn’s multiple comparison adjustment (**j**,**k**).

We therefore hypothesized that mtDNA mutations might impede metastatic entry into the blood. The migratory potential of cybrids was examined under various glucose concentrations that correspond to high (25mM), plasma (5mM) and tumor interstitial fluid (TIF) (1mM) levels^36^. Under conditions of TIF glucose availability, continuous oxygen consumption analysis revealed a significant increase in the oxygen consumption for the WT, ATP6, and ND1 cybrid lines, indicating that TIF conditions stimulate mitochondrial oxidative activity (Extended Data Fig. 8a,b). Notably, these differences were not a consequence of altering cellular viability (Extended Data Fig. 8c-e). Assessment through a scratch-wound assay demonstrated a significant reduction in migration among the pathogenic mtDNA cybrid lines under conditions simulating TIF glucose availability (Fig. 4j, Extended Data Fig 8f,g). Furthermore, a significant decrease in invasion, as evaluated by Boyden transwell assays, was observed in the pathogenic mtDNA cybrid lines under TIF conditions (Fig. 4k, Extended Data Fig. 8h). Pharmacologic inhibition of ETC function with IACS-010759 in the WT cybrid line mirrored the results observed in the mutant mtDNA cybrid lines (Fig. 4j,k). These findings indicate that dysfunctional mtDNA variants limit migration and invasion under metabolic conditions similar to the TIF.

### Evidence of selective pressure during tumor growth favoring functional mtDNA

Although sequence analysis of human tumors has suggested that cancers select for wildtype mitochondrial genomes, our homoplasmic cybrid experiments demonstrate that loss-of-function mitochondrial mutations do not abolish growth of subcutaneous melanoma xenografts. However, these experiments did not address pressures that might restrict the expansion of dysfunctional mtDNA genomes. To probe tumor evolution and potential selective pressures on mtDNA, we generated heteroplasmic cybrid clones by fusing cytoplasts derived from mtDNA dysfunctional donors into the homoplasmic WT cybrid line (Fig. 5a). Immediately following heteroplasmic cybrid fusion, the ATP6, ND1, and CO1 alleles presented as heteroplasmic with their respective wildtype alleles (Extended Data Fig. 9a-c). However, the ND1 and CO1 heteroplasmic models consistently shifted toward increased VAF of the wildtype allele, indicating that these heteroplasmic pathogenic alleles were not stable during clonal expansion (Extended Data Fig. 9d,e). Notably, the ATP6 allele maintained heteroplasmy in culture, as assessed by ddPCR analysis (Fig. 5b). We chose four heteroplasmic ATP6/WT clones for further analysis (Fig. 5b, red arrows). Single-cell analysis of the ATP6 heteroplasmic frequencies in individual clones demonstrated that these clonal lines contained a distribution of single cells with allelic frequencies centered around the calculated allelic frequencies from bulk analysis (Fig. 5c). We noted that the cybrid lines with a higher ATP6 allelic frequency (∼50%, clone 1 and clone 2) exhibited a lower oxygen consumption rate than clones with a lower ATP6 allelic frequency (∼30%, clone 3 and clone 4) (Fig. 5d).

**Figure 5.**
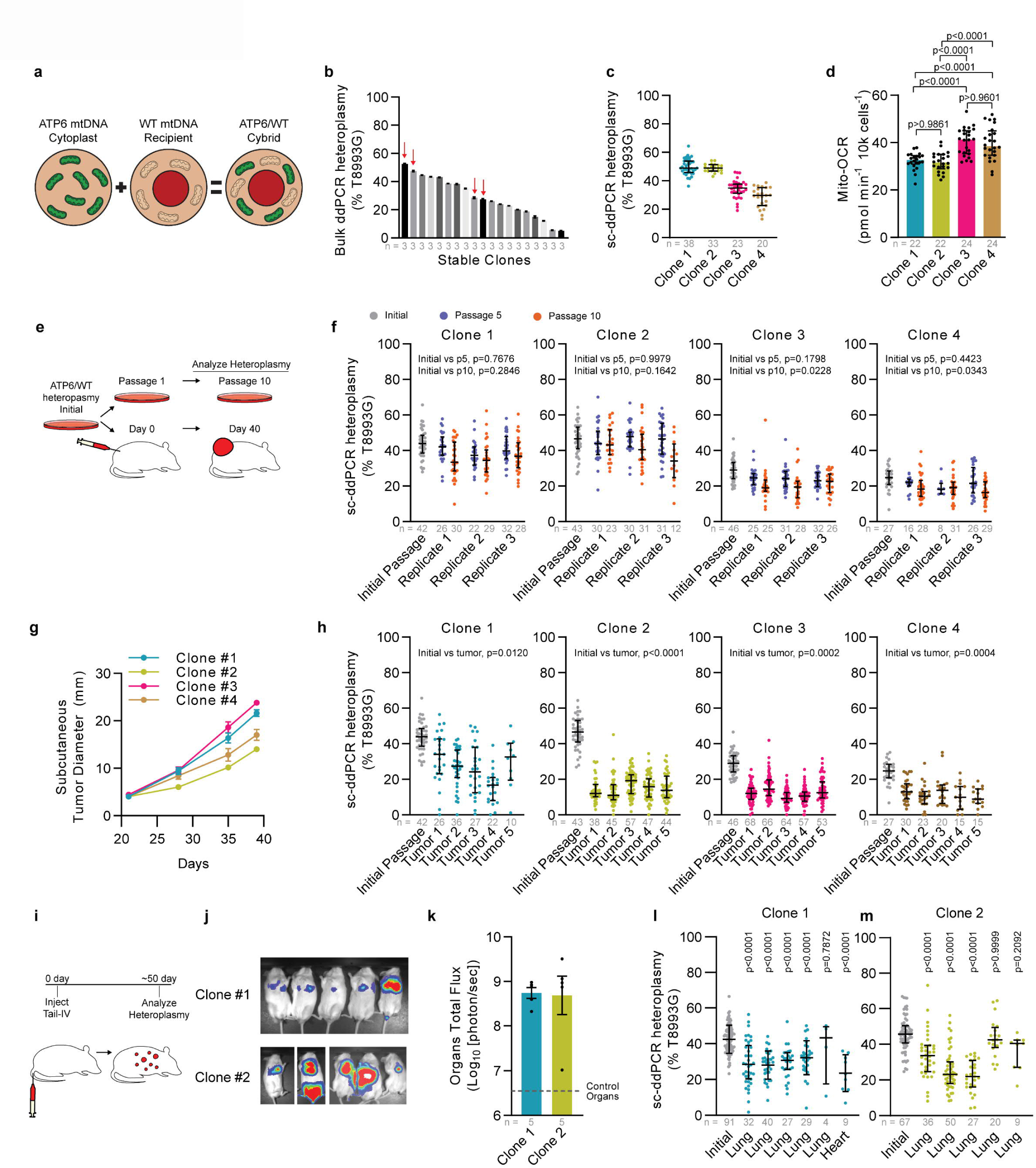
Purifying intracellular mtDNA selection is a feature of melanoma growth. **a**, Overview of ATP6/WT heteroplasmic cybrid generation. **b**, Bulk ddPCR analysis of heteroplasmic frequency at mt.T8993G for ATP6/WT cybrids after clonal line establishment. Clones selected for experiments are indicated with red arrows. **c**, Single cell ddPCR (sc-ddPCR) analysis of heteroplasmy at mt.T8993G for selected ATP6/WT cybrid clones. **d**, Mitochondrial oxygen consumption rate of ATP6/WT heteroplasmic clones. **e**, Illustration of heteroplasmic selection experiment. Initial cells were passaged for extended *in vitro* culture or xenografted for subcutaneous tumors in NSG mice. **f**, Single cell ddPCR analysis of heteroplasmy at mt.T8993G for ATP6/WT heteroplasmic clones at passage 5 (p5) and passage 10 (p10) of *in vitro* culture. Three replicates were independently passaged and analyzed for each clone. **g**, Subcutaneous tumor diameter over time after xenograft of 100 cells from heteroplasmic ATP6/WT clones. **h**, Single cell ddPCR analysis of heteroplasmy at mt.T8993G for ATP6/WT heteroplasmic clones of subcutaneous xenograft of 100 cells following tumor growth. **i**, Illustration of heteroplasmic selection assay following tail-vein IV injection of ATP6/WT heteroplasmic clones. P values reflect comparisons with the initial passage. **j**, Bioluminescence imaging of live mice following intravenous injection of 1,000 cybrid cells. **k**, Quantification of organ bioluminescence total flux following intravenous injections. **l**,**m**, Single cell ddPCR analysis of heteroplasmy at mt.T8993G for ATP6/WT heteroplasmic clones tumor nodules following intravenous injection. The number of samples analyzed per treatment is indicated. Data are mean ± s.e.m. (**b**,**g**), median ± interquartile range (**c**,**d**,**f**,**h**,**k**,**l**). Statistical significance was assessed one-way ANOVA with Tukey’s multiple comparison adjustment (**d**), nested one-way ANOVA with Dunn’s multiple comparison adjustment (**f**,**h**), one-way ANOVA with Dunn’s multiple comparison adjustment (**l**), and non-parametric Kruskal-Wallis test with Dunn’s multiple comparison adjustment (**m**).

These four heteroplasmic ATP6/WT clones, two at ∼50% ATP6 VAF and two at ∼30% ATP6 VAF, were used for concurrent passage in culture and subcutaneous xenografting (Fig. 5e). After subcutaneous xenograft of 100 cells, all clones reached maximal tumor size in ∼40 days (Fig. 5g). In culture, there were minimal shifts in the single cell allelic frequency in these four clones (Fig. 5f). In contrast, we observed that subcutaneous tumors consistently shifted toward increased VAF of the wildtype allele (Fig. 5h). We observed similar results following subcutaneous injection of 10,000 cells per mouse (Extended Data Fig. 10a-e). Further, intravenous injection of heteroplasmic cell lines also shifted toward the wildtype allele in metastatic nodules of visceral organs (Fig. 5i-l). These results indicate that A375 melanoma growth exhibits selection for wildtype mitochondrial genomes when implanted in mice, irrespective of growth in subcutaneous or visceral space.

## Discussion

We identified that isogenic melanoma cybrids transplanted with dysfunctional mitochondrial genomes are capable of sustaining tumor proliferation. Interestingly, the ND1 and CO1 mutant mtDNA cybrid lines displayed a significant reduction in spontaneous metastatic dissemination and all dysfunctional cybrid lines exhibited a reduction in circulating melanoma cells in the blood. Correspondingly, short-term treatment of severe metastatic disease with IACS-010759 ablated the abundance of melanoma cells within the blood of melanoma PDX UT10. In contrast, when mtDNA mutant cybrid lines were delivered through direct intravenous injections, bypassing the process of metastatic circulation entry, they resulted in uniform metastatic seeding regardless of mtDNA mutational status. These results suggest that ETC dysfunctional subcutaneous tumors fail to disseminate in the blood. Moreover, mutant mtDNA cybrid lines exhibited decreased migration and invasion, particularly at the low glucose availabilities similar to the TIF. Therefore, we propose that limited circulation entry functions as a metastatic bottleneck amidst mtDNA dysfunction.

Genomic sequencing analyses of human tumors have revealed that a small number of tumor types (colorectal/renal/thyroid) exhibit enrichment for loss-of-function mtDNA mutations^5-9^. These tumors can exhibit a striking proliferation of mitochondria, resulting in an oncocytic appearance^10^. However, we found that dysfunctional mitochondrial genomes were not sufficient to impart oncocytic features in human melanoma, suggesting tissue specificity and/or alternative alterations drive oncocytoma mitochondrial phenotypes. Further, analysis from pan-cancer mtDNA sequencing studies have suggested that human tumors select against dysfunctional mitochondrial genomes, and here our results provide the first direct experimental verification of this selective pressure. Inherited mitochondrial diseases often manifest in a state of heteroplasmy; thus, insights into the mechanisms that drive tumor selection for functional mtDNA could potentially unveil innovative treatment strategies for mitochondrial disease.

Prior to this report, the impact of complete loss of complex IV function on tumor progression was unknown. The mt.6692del CO1 mutation in this paper was derived from a human melanoma patient derived xenograft model (M405)^27^ and has also been reported in human colonic crypts^37,38^, myopathy^39^, peripheral blood of a breast cancer patient^40^, and within the PCAWG/TCGA dataset for the following cancer types: bone, breast cancer, prostatic adenocarcinoma, esophageal adenocarcinoma, renal cell carcinoma, glioma, hepatobiliary cancer, and non-small cell lung cancer^5,6^. We previously reported metabolic tracing in the mt.6692del M405 PDX model and demonstrated a lack of TCA cycle metabolic activity, as well as minimal metabolic perturbations by treatment with IACS-010759^27^. Here, we build on the effects of mt.6692del (CO1 mutant) and demonstrate that these tumors histologically do not present regions of tumor necrosis or hypoxia, yet exhibit a high proportion of discohesive regions. These results indicate that mitochondrial respiration can contribute to tumor necrotic processes, and future studies expanding to more cancer types will be needed to establish the relationship between severe mtDNA impairment and necrosis.

Lastly, the analyzed partial loss of function cybrid lines, ATP6 and ND1, were derived from well-characterized human mitochondrial disease models. The influence of mitochondrial disease on cancer progression is largely uncharacterized, but will grow in importance as patient survival improves. Within the scope of the studied alleles, these findings suggest that mitochondrial disease may not preclude melanoma development but rather attenuate its severity. Similar studies in other cancer types will be needed to establish the generality of these results.

## Methods

### Experimental models

Immortalized human melanoma cell line A375 (CRL-1619) was obtained from ATCC. Melanoma patient derived xenograft model UT10 was obtained with informed consent according to protocols approved by the Institutional Review Board of the University of Texas Southwestern Medical Center (IRB approval 102010-051). Melanoma specimens used in the manuscript are available, either commercially or from the authors, through there are restrictions imposed by Institutional Review Board requirements and institutional policy on sharing of material from patients.

Immortalized cells were maintained in complete media (DMEM supplemented with 10% fetal bovine serum (FBS), 2 mM L-glutamine, 1 mM sodium pyruvate, 100 µM uridine, and 1% penicillin/streptomycin). For detached experiments, ultra-low attachment surface plates were used (Corning Costar Ultra-Low Attachment Microplates). All cells used in this study were cultivated at 37 °C with 5% CO_2_. Patient derived xenograft UT10 model was maintained in NOD.CB17-Prkdc^scid^ Il2rg^tm1Wjl^/SzJ (NSG) mice (Jackson Laboratories) (details below). Cell culture lines were confirmed as mycoplasma negative using Universal Mycoplasma Detection Kit (ATCC 30-1012K). Cell lines were authenticated through small-tandem repeat analysis by the UT Southwestern McDermott Center Sequencing Core facility.

### Mouse studies and xenograft assays

All mouse experiments complied with all relevant ethical regulations and were performed according to protocols approved by the Institutional Animal Care and Use Committee at the University of Texas Southwestern Medical Center (protocol 2016–101360). For transplantation, cell culture models were trypsin (T4049, Sigma-Aldrich) digested for 5 minutes at 37°C to dissociate from adherent cultures followed by room temperature centrifugation at 200g for 3 minutes. Cells were resuspended, at desired cell count for injection, in staining media (L15 medium containing 1 mg/ml BSA, 1% penicillin/streptomycin and 10 mM HEPES [pH7.4]) with 25% high protein Matrigel (354248; BD Biosciences). PDX single cell suspensions were obtained by mechanically dissociated (12-141-363, Fisher Scientific) on ice followed by enzymatic digestion in 200 U/ml collagenase IV (Worthington), DNase (50 U/ml), and 5 mM CaCl_2_ for 20 minutes at 37°C. Cells were filtered through 40 µm cell strainer to remove clumps, pelleted at 200g for 5 minutes at 4°C, and resuspended in staining media. Subcutaneous injections were performed in the right flank of NSG mice in a final volume of 50 μl. Estimated three-dimensional subcutaneous tumor volume was calculated using the formula: [Length1 × (Length2^2^)]/2. Four to 8-week-old male and female NSG mice were transplanted with 100 or 10,000 melanoma cells subcutaneously as indicated. Intravenous injections were done via tail vein injection into NSG mice with 1,000 melanoma cells in 100 μl of staining media. Mouse cages were randomized between treatments (mice within the same cage received the same treatment). Subcutaneous tumor diameters were measured weekly with calipers until any tumor in the mouse cage reached 2.5 cm in its largest diameter, in agreement with the approved animal protocol. At that point, all mice in the cage were euthanized and spontaneous metastasis was evaluated by gross inspection of visceral organs for macrometastases and bioluminescence imaging of visceral organs to quantify metastatic disease burden (see details below).

For short-term treatment with IACS-010759 (Chemietek), when subcutaneous tumors reached greater than 2.0 cm in diameter, the mice were administered IACS-010759 or control solution by oral gavage daily for 5 days [10 mg/kg body mass in 100 µl of 0.5% methylcellulose and 4% DMSO, adapted from ^27,30^]. On the fifth day and two hours following final oral gavage administration, mice were euthanized and spontaneous metastasis was evaluated.

### Bioluminescence imaging

Metastatic disease burden was monitored using bioluminescence imaging (all melanomas were tagged with a bicistronic lentiviral (FUW lentiviral expression construct) carrying dsRed2 and luciferase (dsRed2-P2A-Luc)). Five minutes before performing luminescence imaging, mice were injected intraperitoneally with 100 μl of PBS containing d-luciferin monopotassium salt (40 mg/ml) (Biosynth) and mice were anaesthetized with isoflurane 2 minutes prior to imaging. All mice were imaged using an IVIS Imaging System 200 Series (Caliper Life Sciences) with Living Image software. Upon completion of whole-body imaging, mice were euthanized and individual organs were surgically removed and imaged. The exposure time ranged from 10 to 60 s, depending upon the maximum signal intensity. To measure the background luminescence, a negative control mouse not transplanted with melanoma cells was imaged. The bioluminescence signal (total photon flux) was quantified with ‘region of interest’ measurement tools in Living Image (Perkin Elmer) software. Metastatic disease burden was calculated as observed total photon flux across all organs in xenografted mice.

### Cell labeling and flow cytometry for circulating melanoma cells

For analysis of circulating melanoma cells, blood was collected from mice by cardiac puncture with a syringe pretreated with citrate-dextrose solution (Sigma) when subcutaneous tumors reached greater than 2.0 cm in diameter. Red blood cells were sedimented using Ficoll, according to the manufacturer’s instructions (Ficoll Paque Plus, GE Healthcare). Remaining cells were washed with HBSS (Invitrogen) prior to antibody staining and flow cytometry. Melanoma cells were identified via flow cytometry as previously described^28,29^. All antibody staining was performed on ice for 20 minutes. Cells were stained with directly conjugated antibodies against mouse CD45 (violetFluor 450, eBiosciences), mouse CD31 (390-eFluor450, Biolegend), mouse Ter119 (eFluor450, eBiosciences) and human HLA-A, B, C (G46–2.6-FITC, BD Biosciences). Cells were washed with staining medium and re-suspended in staining media supplemented with 4’,6-diamidino-2-phenylindole (DAPI; 1 μg/ml; Sigma) to eliminate dead cells from analyses. Human melanoma cells were isolated as events positive for HLA and negative for mouse endothelial and hematopoietic markers.

### Extracellular flux assay

Cells were seeded in Seahorse XFe96 cell culture plates overnight in complete growth media. The next day, cells were washed twice with 200 µl assay medium (DMEM (5030, Sigma-Aldrich) supplemented with 10 mM glucose, 2 mM L-glutamine, 1 mM sodium pyruvate, and 1% penicillin/streptomycin). Subsequently, 150 µl of the assay medium was added to each well. The cells were then placed in a 37°C, CO_2_-free incubator for one hour. Oxygen consumption measurements were conducted on a Seahorse XFe96 instrument, using a three-minute mix and three-minute measure cycle. Three measurements were recorded at baseline and after injecting each compound. Inhibitors were sequentially administered at the specified final concentrations: 2 μM oligomycin, 3 μM CCCP (carbonyl cyanide m-chlorophenyl hydrazone), and 3 μM antimycin A. Data collection and analysis were performed using WAVE software (v.2.4.1.1). Upon completion of the experiment, cells were fixed with formalin, stained with DAPI, and cell counts per well were determined using a Celigo imaging cytometer (Nexcelom Bioscience, 5.1.0.0). Mitochondrial OCR (oxygen consumption rate) was calculated as follows: basal (pre-oligomycin) OCR – baseline (post-antimycin) OCR. Negative Mito-OCR values were replaced with zero. OCR values were normalized based on cell count per well.

### Continuous oxygen consumption assay

A total of 30,000 cells per 96-well were seeded in complete growth media and cultured overnight to form a monolayer. The next day, cells were washed twice with PBS and replaced with DMEM (5030, Sigma-Aldrich) supplemented with 1 mM or 25 mM glucose, 2 mM L-glutamine, and 1% penicillin/streptomycin. Continuous oxygen consumption was monitored for 6 hours with a Resipher instrument (Lucid Technologies Inc.) and analyzed with Lucid Labs Software. OCR values were normalized based on final cell count per well.

### Quantitative PCR (qPCR) for mtDNA / nDNA

For DNA extraction, cells were lysed by digestion with proteinase K (Fisher BioReagents™ Proteinase K Cat No. BP1700-100) in digestion buffer (20 mM Tris, 100 mM NaCl, 0.5% sodium dodecyl sulfate, 10 mM EDTA, pH 7.6) at 44°C overnight. After digestion, the samples were supplemented with additional NaCl to reach a final concentration of 2 mM, enhancing DNA extraction yield. Cellular debris was pelleted via centrifugation at 14,000g for 10 minutes. Total DNA was isolated from the supernatant via phenol chloroform extraction and ethanol precipitation as previously described^41,42^. For mtDNA/nDNA measurements, 3 ng of total DNA was input, and samples were analyzed with iTAQ Universal SYBR Green Supermix (Bio-Rad, #1725120). qPCR analysis was employed to determine the levels of mtDNA (targeting regions 7773-7929) and the nuclear genome for histone H4c, as previously described^43^.

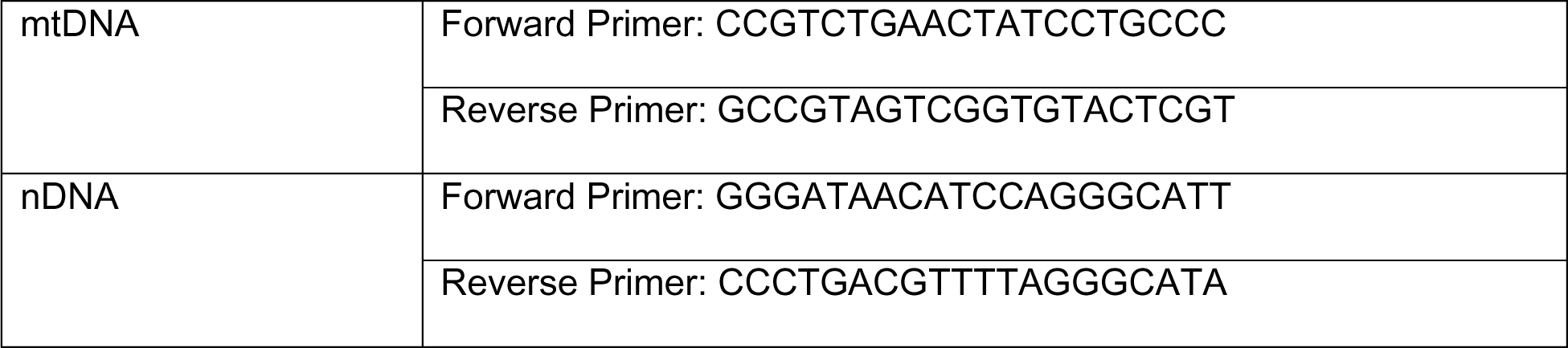

### mtDNA depletion and establishment of stable ρ0 models

Cells were cultured in complete media (DMEM supplemented with 10% fetal bovine serum (FBS), 2 mM L-glutamine, 1 mM sodium pyruvate, 100 µM uridine, and 1% penicillin/streptomycin) and treated with 5 µM or 10 µM dideoxycytidine as indicated. Cells were passaged every two days for two weeks; at which time serial dilution was used to establish single cell colonies. The 5 µM clones demonstrated mtDNA depletion and were selected for cybrid generation (see below).

### Cybrid generation

For cytoplast generation, 1 x 10^6^ - 1 x 10^7^ mitochondrial donors were suspended in 1:1 Percoll (Sigma-Aldrich, 65455-52-9) and cybrid generation buffer (D5030 (Sigma-Aldrich), 25 mM glucose, 2% FBS, 2 mM L-glutamine, 1 mM sodium pyruvate, 100 µM uridine, 1% penicillin/streptomycin, pH 7.2-7.4) supplemented with 10 mg/ml cytochalasin B (Cayman Chemical Company, 14930-96-2), 2000 nM MitoTracker Green FM (Invitrogen, M7514), and 40 µg/ml Hoechst 33342 (Thermo Fisher Scientific, H3570). Cells were then subjected to 39,800g centrifugation for 70 minutes at 37°C. The resulting hazy cellular band located above the percoll cushion was collected (∼5 ml), diluted 10-fold with cybrid generation buffer, and centrifuged at 650g for 10 minutes at room temperature. The cytoplast containing pellet was resuspended in cybrid generation buffer for FACS isolation of cytoplasts defined as MitoTracker Green^+^ and Hoechst^-^.

Recipient cells (either ρ0 cells for homoplasmic cybrids or WT cybrids for heteroplasmic cybrids) were pre-stained with 100 nM SYTO59 (Invitrogen, S11341) for 30 minutes in complete media at 5% CO_2_ and 37°C. Cells were washed 3x with PBS, trypsin digested, centrifuged at 300g for 5 minutes at room temperature, and resuspended in cybrid generation buffer. Cytoplasts isolated via FACS were mixed with recipient cells in a 1:1 ratio. A portion of the mixture (1/10 total volume) was set aside as a non-fusion control for FACS gating. The cell/cytoplast mixture was centrifuged at 300g for 5 minutes at room temperature and all supernatant was removed. The pellet was softly resuspended and mixed with 100 µl of polyethylene glycol Hybri-Max (PEG) (Sigma-Aldrich, P7306) for 1 minute at room temperature. Slowly and with constant mixing, 37°C cybrid generation buffer was added at a rate of 100 µl the first minute, 200 µl the second minute, 300 µl the third minute, and so on up to the seventh minute. Next, over the course of two minutes, 10 mL of cybrid buffer was added to the cells and the post-fusion cell suspension was incubated for 10 minutes at 37°C. The cells were centrifuged at 200g for 7 minutes and resuspended in cybrid generation buffer for FACS isolation of cybrid cells. Successfully fused cybrid cells were defined as MitoTracker Green^+^, Hoechst^-^, and SYTO59^+^. Cybrids were directly sorted into complete media and plated for cell culture.

### Single cell digital droplet PCR quantification of mtDNA

Single cell digital droplet PCR analysis was performed as previously described^44^ with the following exceptions: Single melanoma cells were sorted into cell lysis solution (Proteinase K (Fisher BioReagents™ Proteinase K Cat No. BP1700-100) in 10% [v/v] NP40 (Thermo Fisher, FNN0021), 4.5% [v/v] TNES (50 mM Tris, 0.4 M NaCl, 100 mM EDTA, 0.5% SDS)). Proteinase K digestion was performed at 50°C for 30 minutes, followed by inactivation at 100°C for 10 minutes, then cooled to 12°C. Next, the region of interest was minimally amplified through a 7-cycle PCR (Phusion® High-Fidelity DNA Polymerase, NEB) surrounding the mtDNA regions for ddPCR detection. Following amplification, 15% of the limited PCR product was input for ddPCR probe analysis (Bio-Rad, QX100 droplet reader) following manufacturer instructions. ddPCR data analysis was performed in QuantaSoft Analysis Pro version 1.0.596.

To establish a standard curve, regions of mtDNA were cloned from donor cell lines into p-GEM-T Easy vector (Promega, A1360). Samples were then diluted to specific concentrations and analyzed to evaluate probe specificity.

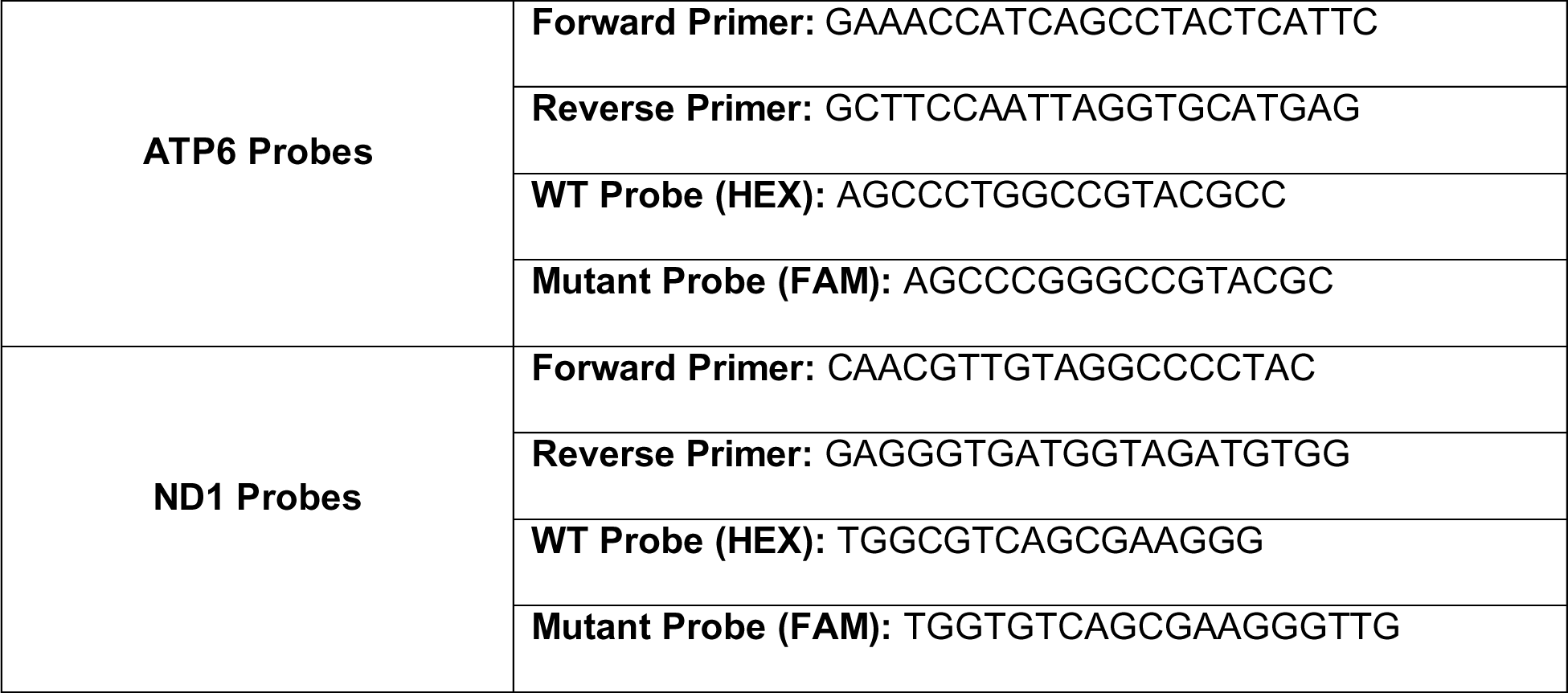

### Histology

Tumors were dissected, fixed in 10% neutral buffered formalin for 48 hours, and paraffin embedded. For pimonidazole staining, three hours prior to collection, mice received an intraperitoneal injection of 60 mg/kg pimonidazole dissolved in saline. For pimonidazole detection 4.3.11.3 mouse FITC-Mab (Hypoxiaprobe, 1:100) was used in combination with mouse on mouse (M.O.M.) blocking IgG reagent (Vector, MKB-2213-1). The following antibodies were used for Ki67 staining: Ki67 (ab15580, Abcam, 1:100) and anti-Rabbit IgG-biotin conjugated (Vector Laboratories, BA-1000-1.5, 1:500). 5 µm serial sections were used for hematoxylin and eosin (H&E) staining, Ki67 staining, and pimonidazole staining. Slides were scanned at 40X using a Nanozoomer 2.0HT (Hamamatsu) at the UTSW Whole Brain Microscopy Facility. QuPath^45^ was used to quantify all histological sections. Quantification of H&E structural regions was performed through training QuPath pixel detection tool on annotated regions. Ki67 stained nuclei were detected using Positive Cell Detection tool in QuPath. Pimonidazole positive regions were detected through applying a threshold in ImageJ^46^.

### Metabolic glucose and lactate consumption and excretion assay

Cells were cultured in adherent culture conditions or detached conditions (Fisher Scientific, 07-200-601) for 24 hours in DMEM supplemented with 10% FBS, 2 mM L-glutamine, and 1% penicillin/streptomycin. Glucose and lactate were measured in culture medium using an automated electrochemical analyzer (BioProfile Basic-4 analyzer, NOVA).

### Migration and invasion assays

Wound healing migration assay was performed as previously described^47^. Briefly, cells were seeded to form a monolayer and subjected to overnight serum starvation. Scratches were created with a sterile p200 tip and wells were washed with PBS to remove detached cells. DMEM (no glucose) supplemented with 10% FBS, 2 mM L-glutamine, 1% penicillin/streptomycin, the appropriate concentration of glucose (1 mM, 5 mM, or 25 mM), and 100 nM IACS-010759 or DMSO vehicle were added to wells. To prevent consumption of all available glucose at lower availabilities, the media was replaced every 8 hours. Images were taken at 0 hour and 24 hour time points with a Celigo image cytometer-4 Channel software version 5.1.0 (Nexcelom Bioscience). The difference in gap length between the 0 hour and 24 hour was reported as distance migrated.

Boyden transwell assay invasion assay was performed as previously described^47^. Briefly, cells were seeded in DMEM (no glucose) supplemented with 1 mM glucose, 2 mM L-glutamine, 1% penicillin/streptomycin, and 100 nM IACS-010759 or DMSO vehicle were added to wells. Following a 6 hour serum starvation, cells were trypsin digested, pelleted at 300g, washed in the appropriate glucose concentration DMEM, and 50,000 cells per well were plated into upper chamber of transwell insert with 8 µm pore size (Celltreat, 2306439). The bottom chamber of the transwell was filled with DMEM (no glucose) supplemented with 10% FBS, 1 mM glucose, 2 mM L-glutamine, 1% penicillin/streptomycin, and 100 nM IACS-010759 or DMSO vehicle. To prevent consumption of all available glucose, the media was replaced every 8 hours. Following 24 hours, the insert was transferred to a PBS wash and a cotton swab (Q-tip) dipped in PBS was used to remove non-migrated cells from the upper chamber. The cells were fixed in 10% buffered formalin for 20 minutes, followed by a 20 minute stain in 0.1% Crystal violet solution (0.1% crystal in 10% ethanol). The inserts were washed 3x with PBS and the upper chamber was cleaned again with a Q-tip. The inserts were allowed to dry for several hours, after which the membrane was cut and imaged with Primovert ZEISS microscope on a ×10 objective. All images were recorded with ZEN 3.1 (Blue ed) software and analyzed with ImageJ.

### Analysis of adherent and detached ROS, viability, and apoptosis

Cells were lifted from adherent passage with PBS supplemented with 1 mM EDTA and 1% FBS. Cells were cultured in DMEM supplemented with 10% FBS, 2 mM L-glutamine, and 1% penicillin/streptomycin for 24 hours in adherent conditions or detached (Ultra-Low Attachment Microplates, Fisher Scientific, 07-200-601) conditions.

For ROS measurements, both conditions were washed with PBS; adherent cells were directly washed with PBS and detached cells were centrifuged at 200g for 3 minutes and resuspended with PBS. Cells were incubated for 30 minutes at 5% CO_2_ and 37°C in staining medium (L15 medium containing bovine serum albumin (1 mg ml−1), 1% penicillin/streptomycin and 10 mM HEPES (pH 7.4)) with 5 µm CellROX Green (Thermo Fisher Scientific, C10444). Cells were washed with staining medium and re-suspended in 4’,6-diamidino-2-phenylindole (DAPI; 1 μg/ml; Sigma) to eliminate dead cells from sorts and analyses. Cells were examined on a FACS Fusion Cell Sorter (Becton Dickinson).

For detached viability, cells were collected and spun down at 200g for 4 minutes. Cell viability was determined by the percentage of cells that were trypan blue positive (Sigma). For apoptosis analysis, the Apo-Direct TUNEL assay kit (Millipore Sigma) was used following manufacturer protocol, and cells were examined on a FACS Fusion Cell Sorter (Becton Dickinson). For adherent monolayer viability, cells were incubated with 1 μg/ml DAPI for 20 minutes at 5% CO_2_ and 37°C. Following, DAPI positive cell counts were determined with a Celigo image cytometer-4 Channel software version 5.1.0 (Nexcelom Bioscience). For total cell counts, cells were fixed with 10% buffered formalin overnight, followed by staining with 1 μg/ml DAPI in PBS for 30 minutes and total cell counts were determined with a Celigo image cytometer-4 Channel software version 5.1.0 (Nexcelom Bioscience).

### Analysis of mitochondrial mass

To assess intracellular mitochondrial mass, adherent cells were washed with PBS and incubated for 30 minutes at 5% CO_2_ and 37°C in staining medium with 20 nM MitoTracker™ Green FM (Thermo Fisher Scientific, M7514). Cells were then washed with staining medium and re-suspended in 1 μg/ml DAPI (Sigma) to eliminate dead cells from sorts and analyses. Cells were examined on a FACS Fusion Cell Sorter (Becton Dickinson).

### Western blot analysis

Tumors were excised and rapidly snap-frozen in liquid nitrogen. Tumor lysates were prepared by mincing tissue in RIPA buffer (Thermo Fisher Scientific, 89901) supplemented with Halt protease and phosphatase inhibitor cocktail (Thermo Fisher Scientific) and maintained in constant agitation for 2 hours at 4°C. For cells, lysis buffer was added to the dish and cells were scrapped on ice before constant agitation for 30 minutes at 4°C. Lysates were spun down at 12,000g at 4°C for 10 minutes. The DC protein assay kit II (BioRad) was used to quantify protein concentrations. Equal amounts of protein (5 µg) were loaded into each lane and separated on 4– 20% polyacrylamide Tris glycine SDS gels (BioRad), then transferred to 0.45 µm PVDF membranes (Millipore Sigma). The membranes were blocked for 1 h at room temperature with 5% milk in TBS supplemented with 0.1% Tween-20 (TBST) and then incubated with primary antibodies overnight at 4 °C. After washing and incubating with horseradish peroxidase conjugated secondary antibody (Cell Signaling Technology), signals were developed using Immobilon Western Chemiluminescent HRP Substrate (Millipore Sigma). Blot data collection was performed using Amersham ImageQuant 800. Blots were sometimes stripped using Restore PLUS stripping buffer (Thermo Fisher) and re-stained with other primary antibodies. The following antibodies were used for western blots:anti-mtCO1 (ab14705, Abcam), anti-ATP8 (26723-1-AP, Proteintech), anti-HSP60 (15282-1-AP, Proteintech), anti-TOMM20 (11802-1-AP, Proteintech), and anti-β-actin (4970, Cell Signaling).

### Statistical analysis

Mice were allocated to experiments randomly and samples processed in an arbitrary order, but formal randomization techniques were not used. Samples sizes were not pre-determined based on statistical power calculations but were based on our experience with these assays. For assays in which variability is commonly high, we typically used n>10. For assays in which variability is commonly low, we typically used n<10. All data representation is indicated in the figure legend of each figure. No blinding or masking of samples was performed. All represented data are unique biological replicates.

Prior to analyzing the statistical significance of differences among treatments, we tested whether the data were normally distributed and whether variance was similar among treatments. To test for normal distribution, we performed the Shapiro–Wilk test. To test if variability significantly differed among treatments we performed F-tests. When the data significantly deviated from normality or variability significantly differed among treatments, we log_2_-transformed the data and tested again for normality and variability. Fold change data were log_2_-transformed. If the transformed data no longer significantly deviated from normality and equal variability, we performed parametric tests on the transformed data. If the transformed data remained significantly deviated from normality or equal variability, we performed non-parametric tests on the non-transformed data. For normally-distributed data, groups were compared using the two-tailed Student’s t-test (for two groups), or one-way ANOVA or two-way ANOVA (>2 groups), followed by Dunnett’s or Tukey’s test for multiple comparisons. For data that was not normally distributed, we used non-parametric testing (Kruskal-Wallis test for multiple groups), followed by Dunn’s multiple comparisons adjustment. All statistical analyses were performed with GraphPad Prism 9.5.1.

## Author Contributions

S.D.S., A.T., and P.M. conceived the project. S.D.S. performed experiments with technical assistance from S.H., V.R., T.W., X.W., C.B.L., S.S.K.V., and C.J.M. Z.K. established and characterized A375 ρ0 clones. S.D.S, Z.Z., and J.G.G. contributed to formal analysis and data curation. A.T., R.J.D., and S.J.M. contributed to the investigation, provided critical resources, and reviewed and edited the manuscript. S.D.S and P.M. wrote the original draft and prepared figures. P.M., R.J.D., S.J.M. provided supervision and acquired funding.

## Acknowledgements

We thank the Moody Foundation Flow Cytometry Facility, the Children’s Research Institute Next Generation Sequencing Facility, the UT Southwestern Histo Pathology Core, the UT Southwestern Whole Brain Microscopy Facility, and the UT Southwestern BioHPC supercomputing facility. We acknowledge the assistance of ChatGPT, which served to improve grammatical accuracy and clarity while leaving the ideation and conceptualization entirely to the authors. We thank M. Mulkey for mouse colony management. The research was supported by the Cancer Prevention and Research Institute of Texas (RP180778 to S.J.M, R.J.M, P.M.), by the National Institutes of Health (1DP2ES030449-01 from NIEHS to P.M., 1R01AR073217-01 from NIAMS to P.M., R35CA22044901 from NCI to R.J.D., and U01CA228608 from NCI to S.J.M.), H.H.M.I funding (to R.J.D. and S.J.M.), and the Moody Medical Research Institute (research grant to P.M.). S.D.S. was supported by the National Science Foundation (GRF 2019281210), A.T. was supported by an Emmy Noether Award from the German Research Foundation (DFG, 467788900) and the Ministry of Culture and Science of the State of North Rhine-Westphalia (NRW-Nachwuchsgruppenprogramm). A.T. holds the Peter Hans Hofschneider of Molecular Medicine endowed professorship by the Stiftung Experimentelle Biomedizin.

## Data and Material Availability

All data and materials are available from the corresponding author upon request.

---

**Extended Data Figure 1.**
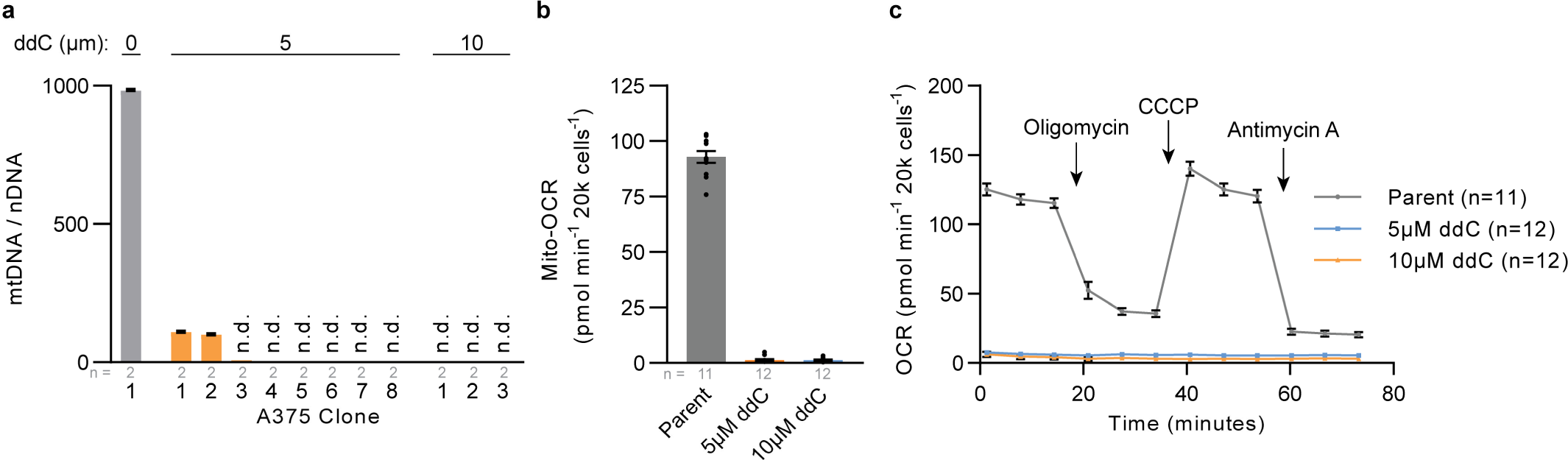
A375 mtDNA depletion and establishment of ρ0 clones. **a**, Mitochondrial genome (mtDNA) to nuclear genome (nDNA) ratios in A375 parental line and serial diluted clones following two week treatment with 5 μM or 10 μM ddC (dideoxycytidine). n.d., mtDNA not detected. **b**,**c**, Mitochondrial oxygen consumption rate (mito-OCR) (**b**) and representative oxygen consumption rates (**c**) for 20,000 cells (per well) of A375 parental line and ddC treated clones. Mitochondrial inhibitors (oligomycin, CCCP, antimycin A) were injected at the indicated timepoints. The number of samples analyzed per treatment is indicated. Data are mean ± s.e.m.

**Extended Data Figure 2.**
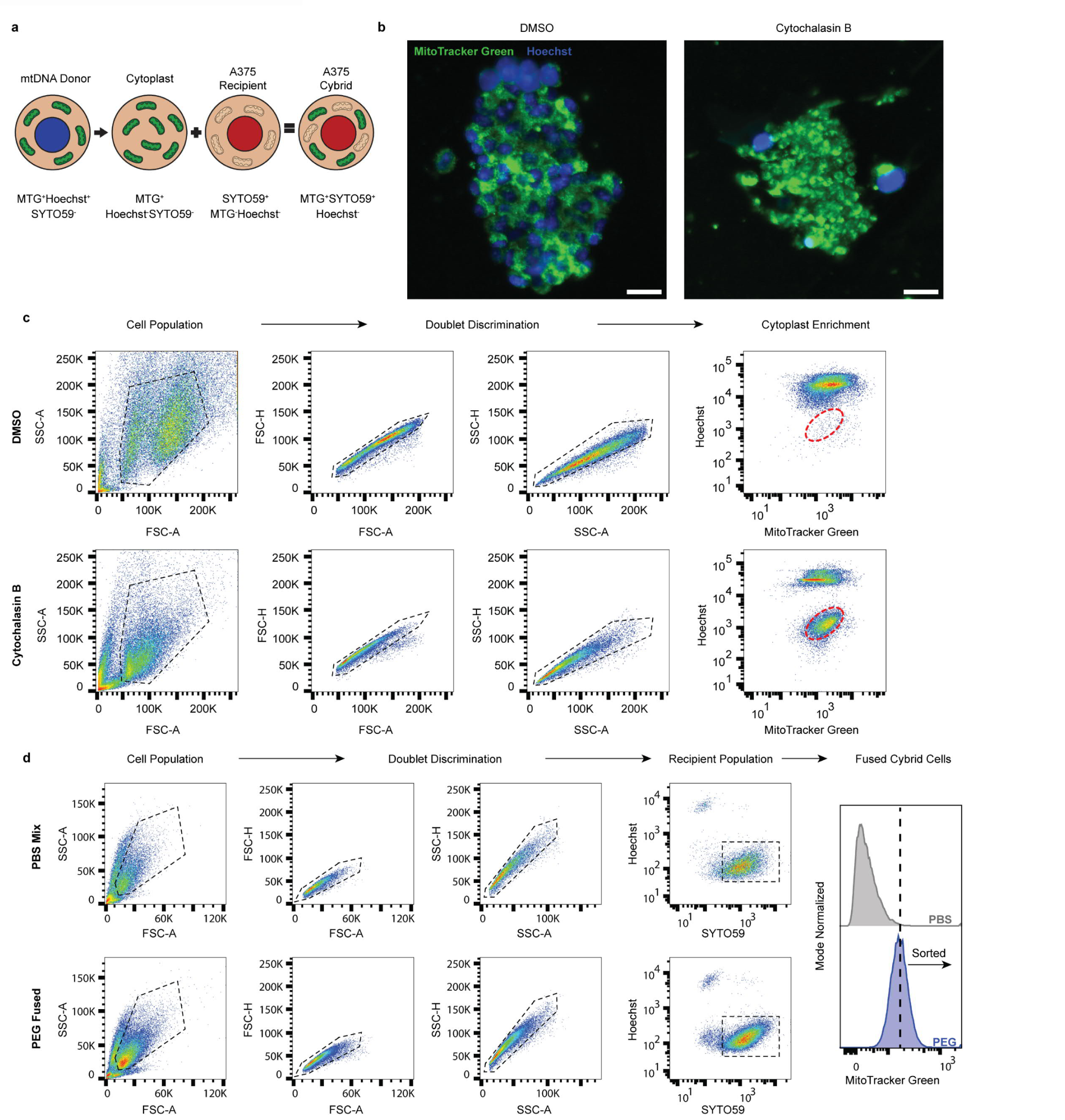
Generation of cybrid cell lines. **a**, Illustration of the cellular fractionation and staining utilized during generation of A375 cybrids. Donor mtDNA lines were stained with MitoTracker Green and Hoechst prior to cytoplast generation. Cytoplasts were generated, enrich via flow cytometry, and then fused with mtDNA depleted A375 ρ0 cells that were pre-stained with SYTO59. Successful cybrid cells were further enriched based on the presence of SYTO59 signal, MitoTracker Green signal, and the absence of Hoechst signal. **b**, Representative epifluorescence images of stained mtDNA donor 143B_nuclear_ Wildtype_mtDNA_ cells following treatment with DMSO vehicle (left) and cytochalasin B (right) and centrifugation to generate cytoplasts. Scale bars, 25 μm. **c**, Enrichment of cytoplasts following differential centrifugation with treatment of DMSO (top) or cytochalasin B (bottom). Cell gating populations are displayed with black dashed lines and sorted cytoplast population is shown with red dashed lines. **d**, Flow cytometric enrichment of PBS mixed (top) and PEG fused cybrid cells (bottom). Cell gating populations are displayed with black dashed lines. Fusion positive populations were sorted for an enrichment of MitoTracker Green in the PEG fused cybrid cells relative to the PBS mixed cells.

**Extended Data Figure 3.**
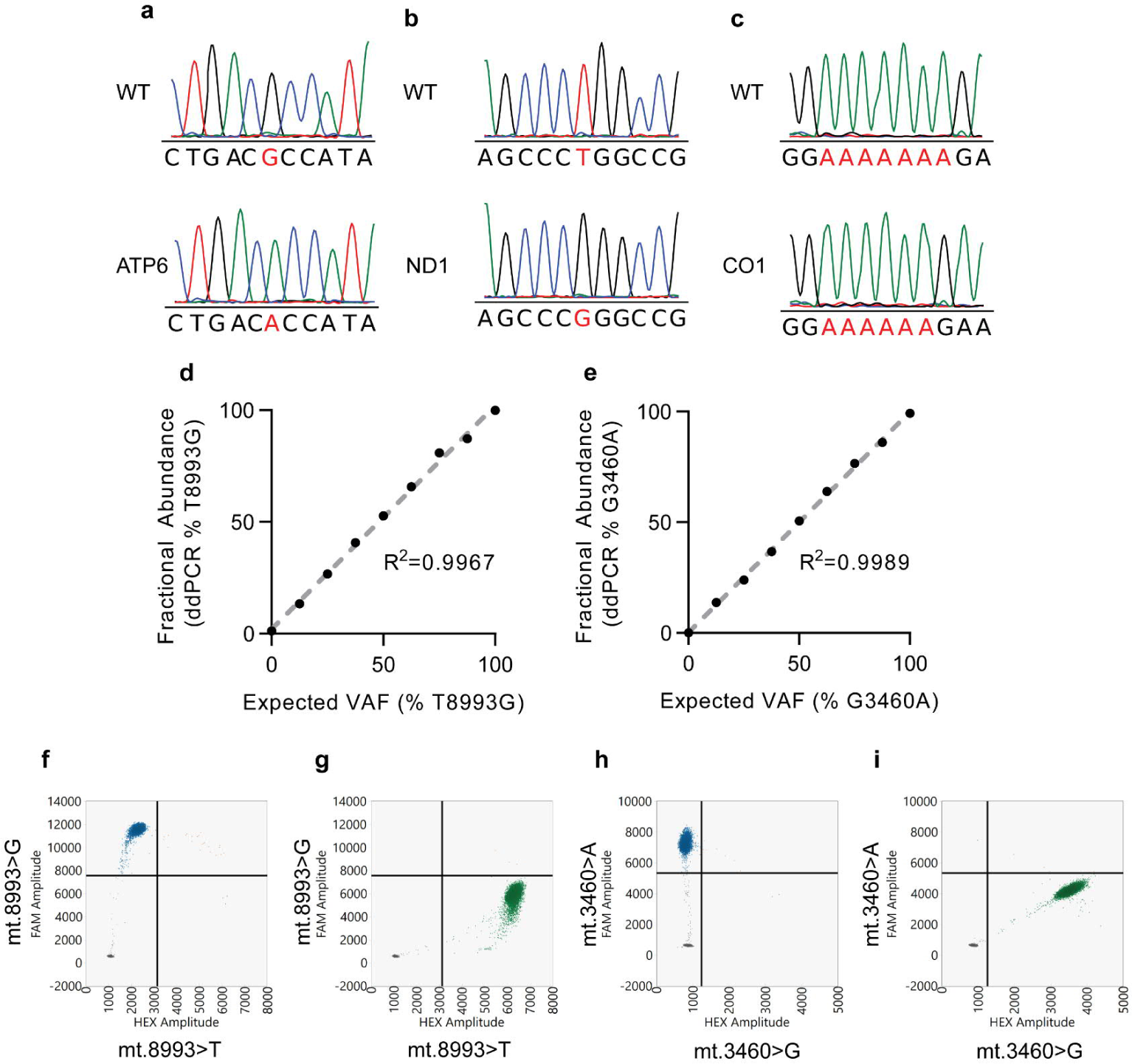
Genetic validation of homoplasmic A375 cybrid lines. **a**,**b**,**c**, Representative Sanger sequencing of mtDNA regions surrounding mt.3460 (**a**), mt.8993 (**b**), and mt.6692 (**c**) for wildtype cybrid line (top) and indicated variant cybrid lines (bottom). Pathogenic variant allelic locations are highlighted in red. **d**,**e**, Standard curve for ddPCR probes specific to mt.8993 T>G (**d**) and mt.3460 G>A (**e**) constructed from purified plasmids for each variant. The coefficient of determination (R^2^) is provided for each curve. **f**,**g**, Representative ddPCR 2-dimensional plot with probes specific to the ATP6 T8993G in homoplasmic ATP6 cybrids (**f**) and wildtype cybrids (**g**). **h**,**i**, Representative ddPCR 2-dimensional plot with probes specific to the ND1 G3460A in homoplasmic ND1 cybrids (**h**) and wildtype cybrids (**i**).

**Extended Data Figure 4.**
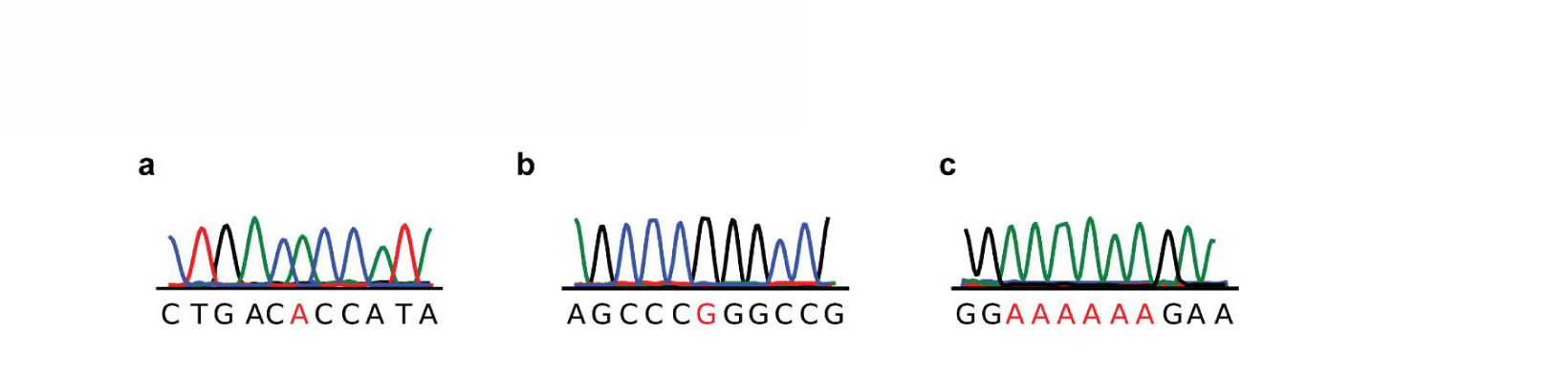
Retention of mtDNA variants after subcutaneous tumor development. **a,b,c,** Representative Sanger sequencing of mtDNA regions surrounding mt.3460 (ND1) (**a**), mt.8993 (ATP6) (**b**), and mt.6692 (CO1) (**c**) from DNA of xenograft subcutaneous tumors for each respective cybrid lines. Pathogenic variant allelic locations are indicated in red.

**Extended Data Figure 5.**
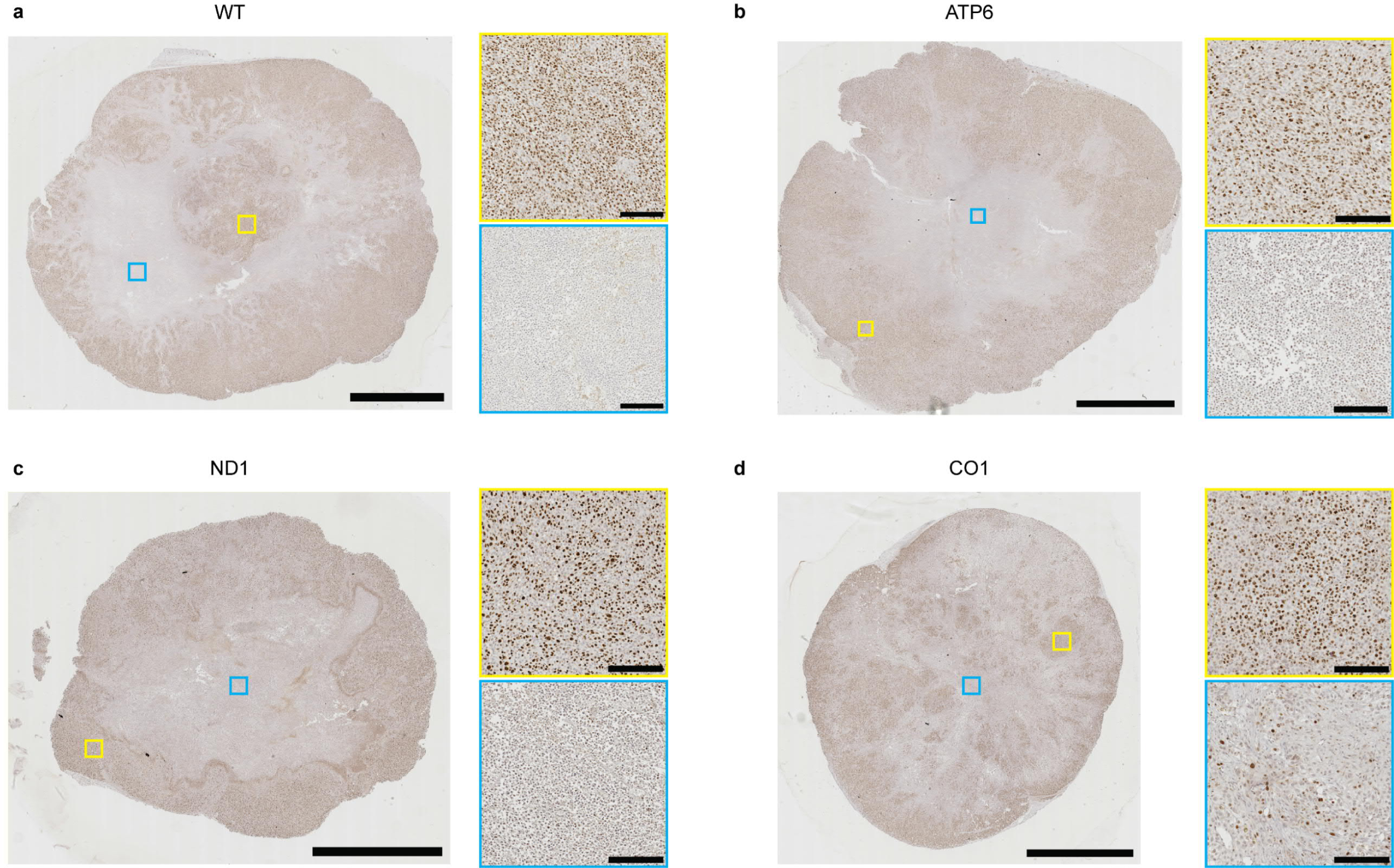
Representative Ki67^+^ staining of subcutaneous cybrid tumors. **a-d**, Representative Ki67 staining images of xenograft subcutaneous tumors of the indicated mtDNA haplotype. Highlighted are proliferative regions with high abundance of Ki67^+^ nuclei (top yellow box) and non-proliferative regions with low abundance of Ki67^+^ nuclei (bottom blue box). Scale bar for full section, 5000 μm. Scale bar for zoomed region, 200 μm.

**Extended Data Figure 6.**
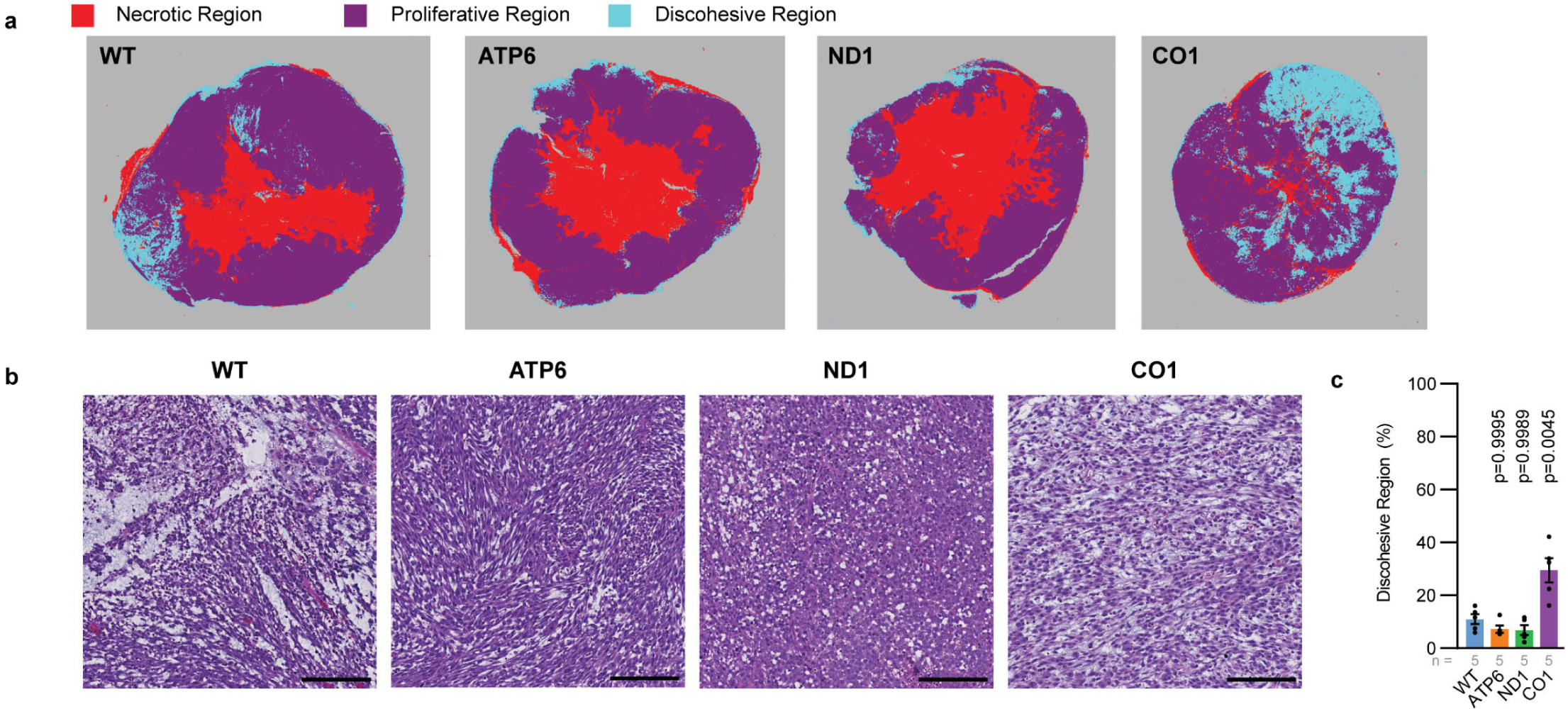
Histological analysis of subcutaneous tumors. **a**, Quantitation of tumor regions from representative H&E sections (Fig 2e) with QuPath Analysis Pixel Classification algorithm. **b**,**c**, Representative regions (**b**) and cross-sectional quantitation (**c**) of discohesive regions in H&E sections of indicated xenograft subcutaneous cybrid tumors. Scale bar, 500 µM. P values indicate comparison with WT group. The number of tumors analyzed per treatment is indicated. Data are mean ± s.e.m. (**c**). Statistical significance was assessed using one-way ANOVA with Dunn’s multiple comparison adjustment (**c**).

**Extended Data Figure 7.**
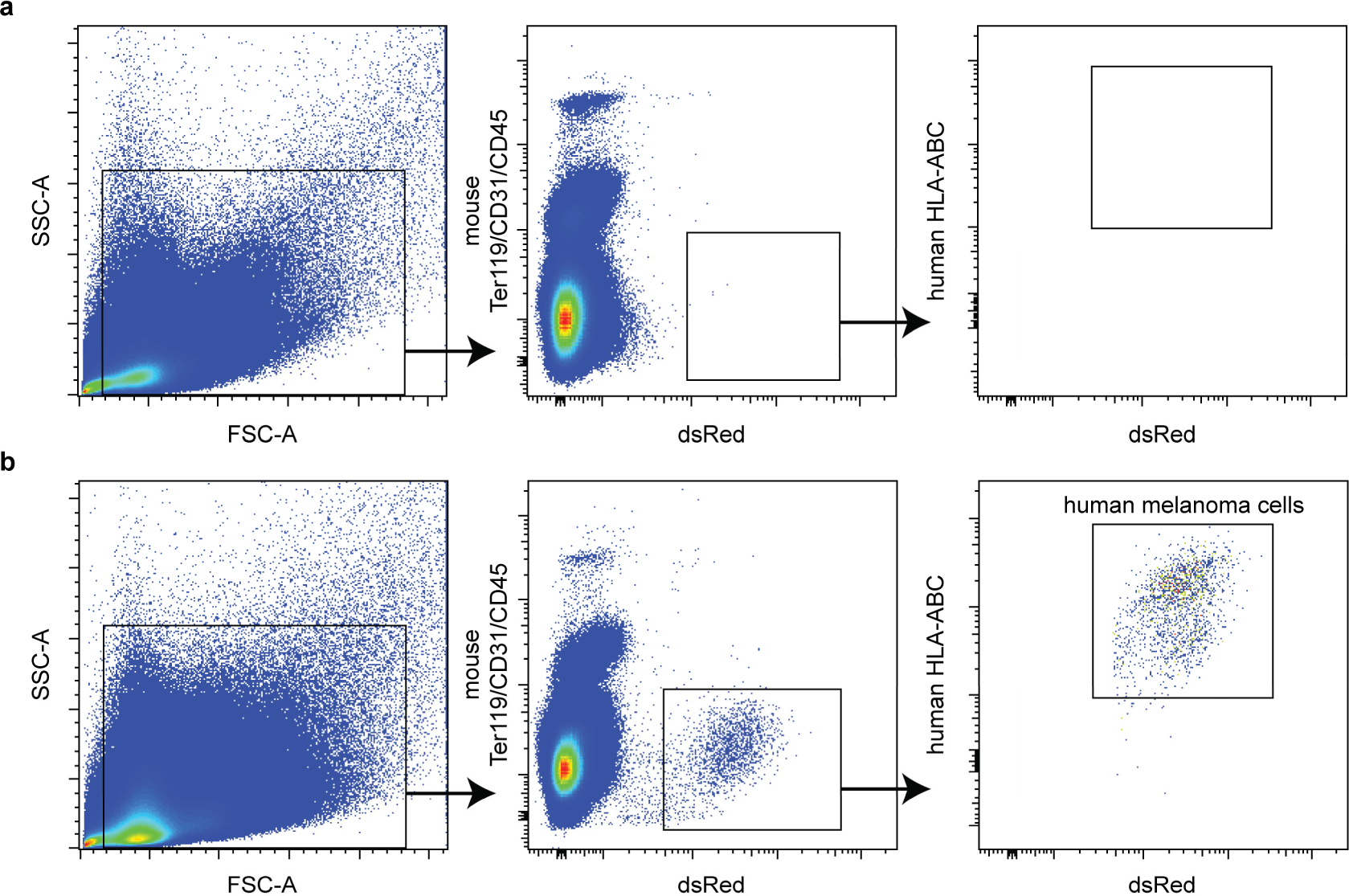
Isolation of circulating human melanoma cells from mouse blood. **a**,**b**, Flow cytometric analysis and gating strategy to identify human melanoma cells in negative control mouse blood (**a**) or blood of xenograft subcutaneous tumor bearing mouse (**b**).

**Extended Data Figure 8.**
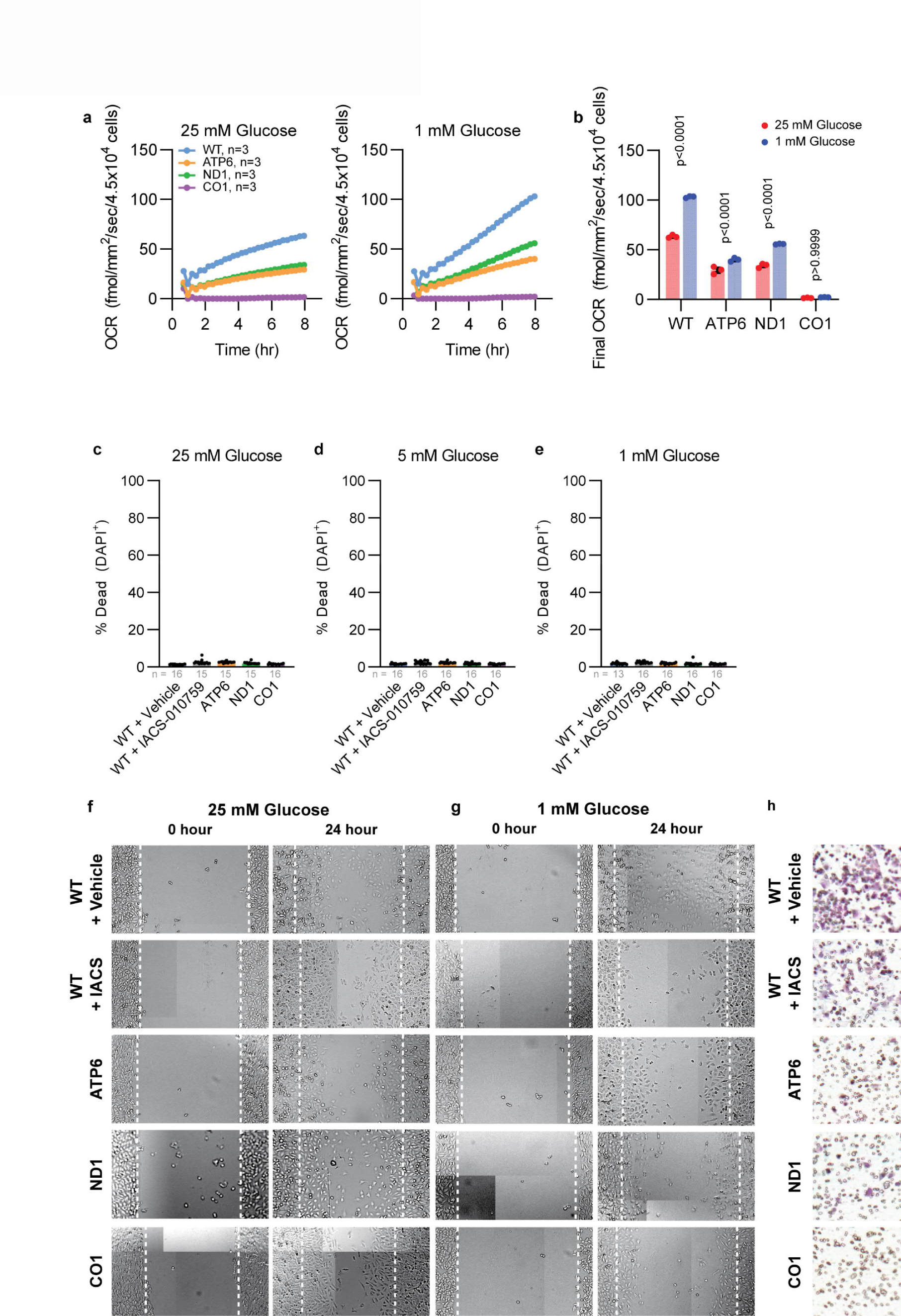
Reduced migration and invasion of dysfunctional cybrid lines at low glucose availability. **a,b,** Continuous (**a**) and final measurement (**b**) of oxygen consumption by a monolayer culture of indicated cybrid lines at 1 mM and 25 mM glucose. P values indicate comparison of 1 mM and 25 mM oxygen consumption. **c-e,** Viability for indicated cybrid lines at 25 mM (**a**), 5 mM (**b**), and 1 mM (**c**) glucose concentrations after 24 hours of culture as a confluent monolayer. **f,g,** Representative wound healing migration images after 24 hours of culture in 25 mM glucose and 1 mM glucose media. Image is composed of individually stitched images with contrast enhanced for viewing purposes. **h,** Representative Boyden transwell migration of cybrid cells after 24 hours of culture in 1 mM glucose media. Image contrast was enhanced for viewing purposes. The number of cells analyzed per treatment is indicated. Data are mean ± standard error of the mean (**a**,**b**) and median ± interquartile range (**c**-**e**). Statistical significance was assessed using two-way ANOVA with Šídák’s multiple comparisons test (**b**).

**Extended Data Figure 9.**
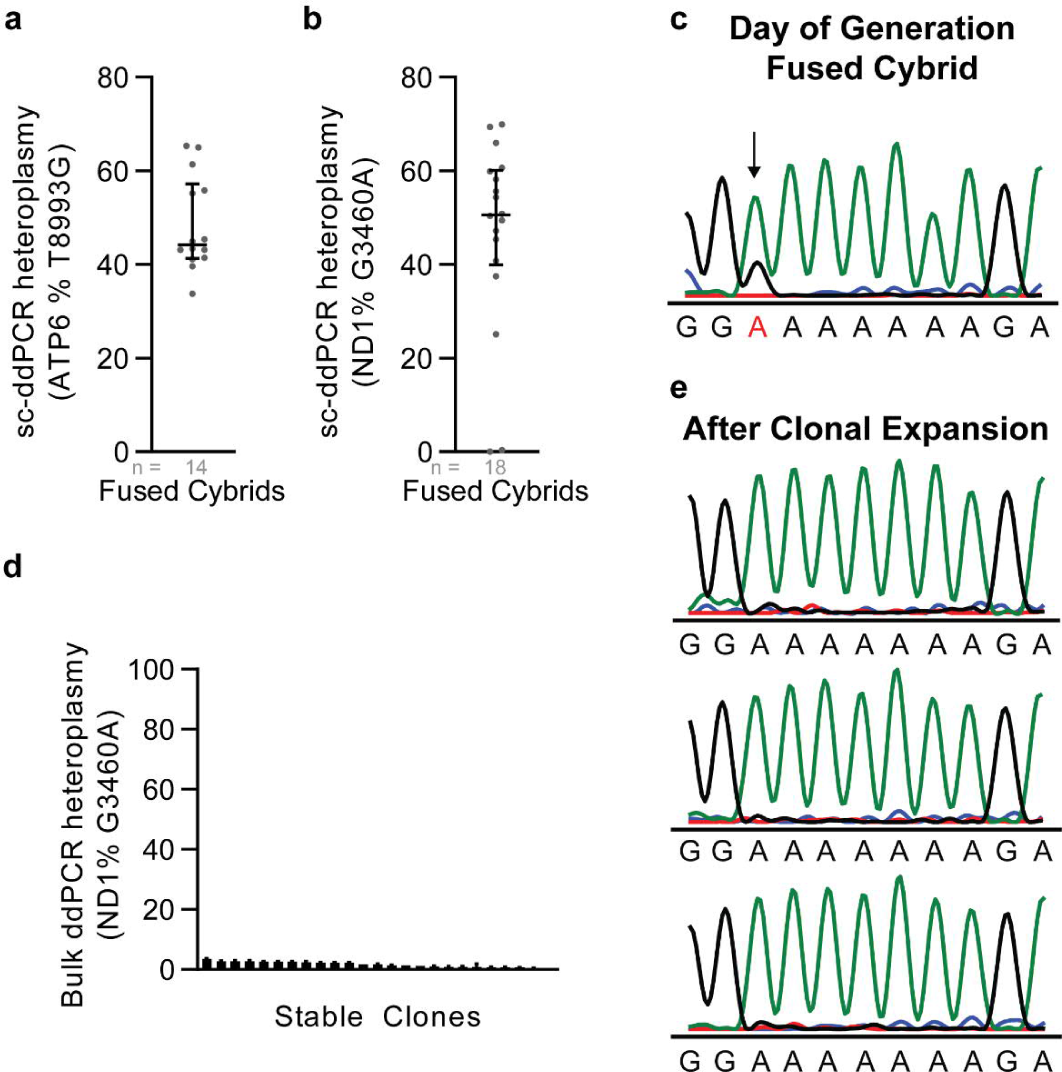
Heteroplasmic ND1 and CO1 alleles are lost after passage in culture. **a**,**b**, Single cell ddPCR (sc-ddPRCR) analysis of heteroplasmy at ATP6 mt.T8993G (**a**) and ND1 mt.G3460A (**b**) directly following cybrid fusion of respective mutant cytoplasts with WT cybrid clones. **c**, Sanger sequencing at CO1 mt.6692del directly following cybrid fusion of CO1 cytoplasts with WT cybrid clones. Heteroplasmic allele is indicated with black arrow and red font. **d**, Bulk ddPCR analysis of heteroplasmic frequency at mt.G3460A for ND1/WT cybrids after clonal line establishment. **e**, Sanger sequencing at CO1 mt.6692del for three representative CO1/WT cybrids after clonal line establishment. Data are median ± interquartile range (a,b) and mean ± SEM (d).

**Extended Data Figure 10.**
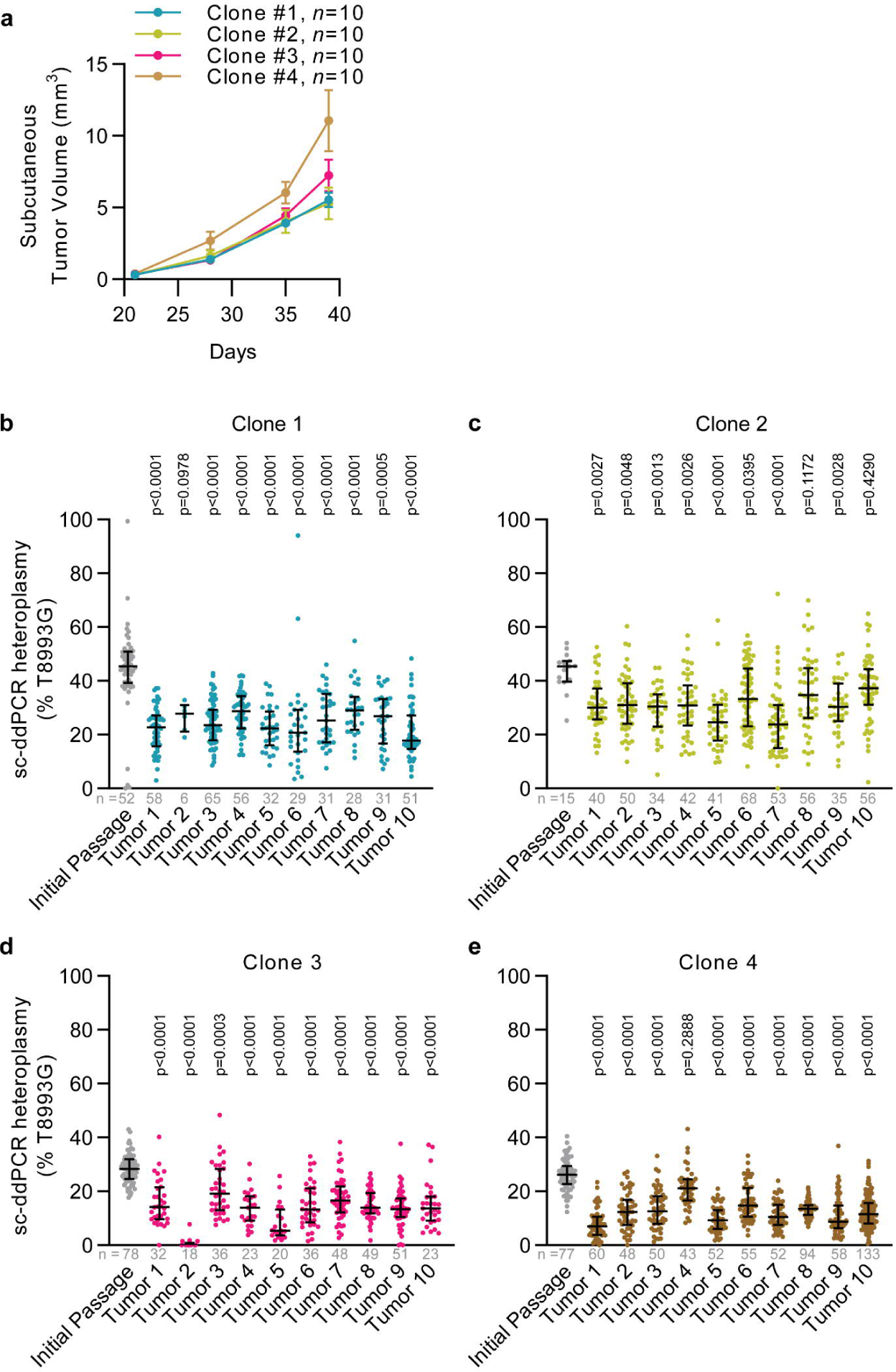
Heteroplasmy assessment of ATP6/WT subcutaneous tumors at increased initial subcutaneous injection cell count. **a**, Subcutaneous tumor volume over time after xenograft of 10,000 cells from heteroplasmic ATP6/WT clones. **b**-**e**, Single cell ddPCR analysis of heteroplasmy at mt.T8993G for ATP6/WT heteroplasmic clones of subcutaneous xenograft of 10,000 cells following tumor growth of indicated clones. P values reflect comparisons with the initial passage. The number of cells analyzed per treatment is indicated. Data are median ± interquartile range (**b**-**e**). Statistical significance was assessed using non-parametric Kruskal-Wallis test with Dunn’s multiple comparison adjustment (**b**-**e**).

